# Adapting C_4_ photosynthesis to atmospheric change and increasing productivity by elevating Rubisco content in Sorghum and Sugarcane

**DOI:** 10.1101/2024.05.02.592081

**Authors:** Coralie E. Salesse-Smith, Noga Adar, Baskaran Kannan, Thaibinhduong Nguyen, Wei Wei, Ming Guo, Zhengxiang Ge, Fredy Altpeter, Tom E. Clemente, Stephen P. Long

## Abstract

Meta-analyses and theory show that with rising atmospheric [CO_2_], Rubisco has become the greatest limitation to light-saturated leaf CO_2_ assimilation rates (*A*_sat_) in C_4_ crops. So would transgenically increasing Rubisco increase *A*_sat_ and result in increased productivity in the field? Here, we successfully overexpressed the Rubisco small subunit (*RbcS*) with Rubisco accumulation factor 1 (*Raf1*) in both sorghum and sugarcane, resulting in significant increases in Rubisco content of 13-25% and up to 90% respectively. *A*_sat_ increased 12-15% and Rubisco enzyme activity ∼40% in three independent transgenic events of both species. Sorghum plants also showed increased speeds of photosynthetic induction and decreased bundle sheath leakiness. These improvements translated into average increases of 15.5% in biomass in field-grown sorghum and a 37-81% increase in greenhouse-grown sugarcane. This suggests a potential opportunity to achieve substantial increases in productivity of this key economically important clade of C_4_ crops, future proofing their value under global atmospheric change.

**Significance Statement:** The world is projected to need a 60% increase in food supply by 2050 (UN), and this must be achieved under conditions of global change without expanding onto yet more land. C_4_ crops, while few in number, account for a large proportion of agricultural productivity. We reason that rising atmospheric [CO_2_] has very recently made Rubisco, the enzyme used for all carbon fixation in plants, the greatest limitation to light saturated photosynthesis in C_4_ crops. We demonstrate that transgenically increasing Rubisco content in sorghum and sugarcane, increases their photosynthetic efficiency and productivity, including in a field trial of sorghum. This shows a means to sustainably increase the productivity of this key group of crops.

## Introduction

While very few of the 161 crop species listed by UN-FAO use C_4_ photosynthesis, their importance and potential in terms of global food, feed and biomass production is exceptional (1). For total global biomass harvested, sugarcane (*Saccharum* hybrids) is number one and maize (*Zea mays* L.) number one for seed harvested (1, 2). Sorghum (*Sorghum bicolor*) is the fourth most important cereal in global production, only behind, maize, rice and wheat, but it is the major cereal of the semi-arid tropics (1). While grown for biomass, the C_4_ crops energycane and Miscanthus show exceptional productivity (3–6). These crops all belong to the monophyletic grass tribe Andropogonae.

The leaves of C_4_ plants contain an outer layer of mesophyll cells surrounding the bundle sheath cells. CO_2_ diffuses from the sub-stomatal cavity into the mesophyll cytoplasm, where it combines with phosphoenolpyruvate (PEP) to form oxaloacetate (OAA), catalyzed by cytoplasmic PEP carboxylase (PEPC). OAA is reduced to malate in the mesophyll chloroplasts and then diffuses to the bundle sheath where it is decarboxylated by NADP^+^-dependent malic enzyme (NADP-ME) creating a high [CO_2_] around Ribulose-1:5 bisphosphate carboxylase/oxygenase (Rubisco). The decarboxylation product, pyruvate, diffuses back to the mesophyll where, catalyzed by Pyruvate Pi dikinase (PPDK), it is phosphorylated to PEP at the expense of 2 ATP, completing the photosynthetic C_4_ cycle (7). This photosynthetic C_4_ cycle is in effect a light energy driven CO_2_ pump that elevates the [CO_2_] around Rubisco to a level that is sufficient to competitively inhibit oxygenation, preventing photorespiration and avoiding the significant losses of photosynthetic efficiency that this causes in C_3_ species (8). Its benefits are such that considerable effort is being placed into engineering the C_4_ pathway into C_3_ crops (9–11). Tests under open-air conditions using Free-Air Concentration Enrichment (FACE) technology, have shown that higher [CO_2_] increases the photosynthetic efficiency, and in turn productivity, of C_3_ crops, but not these C_4_ crops (12–16). Thus, rising atmospheric [CO_2_] could progressively erode the current production advantage of C_4_ plants (17). However, analysis of metabolic and physiological limitations indicate that C_4_ plants could be modified to also gain photosynthetic efficiency at today’s and future atmospheric [CO_2_] (18–20).

Light-saturated leaf CO_2_ assimilation rates (*A*_sat_) in C_4_ plants rise sharply with intercellular CO_2_ concentration (*C*_i_) to a plateau, where *A*_sat_ remains constant with further increase in *C*_i_ (7). In maize crops grown in the field at both ambient and elevated atmospheric CO_2_, this transition was shown to occur at a *C*_i_ of about 120 μmol mol^−1^, above which *A*_sat_ is [CO_2_]-saturated (15). A meta-analysis of 842 measured *A*-*C*_i_ responses in 49 C_4_ species showed the point of CO_2_ saturation is consistent across a broad range of C_4_ species, and unaffected by growth at either sub-ambient or supra-ambient [CO_2_] (19). Because of their lower stomatal conductance (*g*_s_) relative to *A*_sat_, the drawdown of [CO_2_] between the atmosphere (*C*_a_) and intercellular air space (*C*_i_) is greater in C_4_ leaves, resulting in a ratio (*C*_i_/*C*_a_) of about 0.4, compared to 0.7 in C_3_ species.

These ratios in non-water stressed plants are apparently unaffected by the [CO_2_] of the growth environment (12). At today’s *C*_a_ of 428 μmol mol^−1^ (21), *C*_i_ in photosynthesizing C_4_ leaves should be about 170 μmol mol^−1^. Therefore, at current and future atmospheric [CO_2_], C_4_ photosynthesis is [CO_2_] saturated. Historically this was not the case. The average *C*_a_ of the last 40,000 years in which our crop ancestors evolved was 220 μmol mol^−1^ and assuming a *C*_i_/*C*_a_, of 0.4 under these conditions average *C*_i_ would have been 88 μmol mol^−1^, well below the indicated saturation point of about 120 μmol mol^−1^ (12, 22). At this *C*_i_ photosynthesis would be operating on the initial slope of the response of *A* to *C*_i_. The mechanistic steady-state mathematical model of C_4_ photosynthesis shows that this initial slope is determined by the activity of PEPC and mesophyll conductance (*g*_m_); *g*_m_ being the ease with which CO_2_ can diffuse from the sub-stomatal cavity into the mesophyll cytoplasm (7). The plateau however, is limited by the activity of Rubisco, and potentially electron transport (7). Recent non-steady state metabolic modeling and measurements show Rubisco activity also having significant control over the efficiency of photosynthesis in fluctuating light (23, 24). Consistent with these predictions, transgenic knock-down studies showed metabolic control of the initial slope of the *A*_-_*C*_i_ response to reside with PEPC activity and the plateau shared by Rubisco and PPDK, with Rubisco accounting for 70% of the control (25). This indicates that upregulation of Rubsico would increase *A*_sat_ in today’s elevated [CO_2_] atmosphere.

Form I Rubisco, found in plants, is made up from eight small subunits (SS) encoded by the nuclear gene family *RbcS* and eight large subunits (LS) encoded by the chloroplast gene *rbcL*. Previously, transgenic increase in Rubisco content, by overexpression of the Rubisco small-subunit (*RbcS*) and Rubisco accumulation factor 1 (*Raf1*) genes, increased *A*_sat_ and plant growth of maize under optimal and chilling growth conditions (26, 27). However, this was confined to a single hybrid genotype of maize grown in controlled environments. To test the hypothesis that this improvement would be more generally possible across this key economic clade of crops and result in increased productivity in the field and greenhouse, Rubisco was upregulated in both sorghum and sugarcane.

## Results

### Transgenic upregulation of *RbcS* and *Raf1* genes increases Rubisco abundance in sorghum and sugarcane

Two transgenes, each designed to upregulate *RbcS* and *Raf1*, were stably transformed into sorghum and sugarcane (Fig.1a). The construct transformed into sorghum contains constitutive promoters, while the one transformed into sugarcane contains bundle sheath specific promoters as well as C-terminal epitope tags FLAG and HA. For sorghum, 9 transgenic events were received and 7 determined to be single insertion lines. Transgene expression was confirmed by copy number analysis and Western blot (Figure S1a). T1 plants from 7 events were screened for increased CO_2_ assimilation (*A*) and Rubisco activity (*V*_c,max_) using gas exchange (Fig. S1b-g). For sugarcane, 49 transgenic events were received and analyzed via Western blot and a subset of events screened for improved *A* and *V*_c,max_ (Fig. S2). The top three performing events from each species based on gas exchange and protein data were then characterized along with a wild-type (WT), from which the transformants were derived, propagated alongside these. In sorghum, homozygous plants were obtained from each single insertion event (hn1a, zg12a and zg5b) at T2. Commercial sugarcane is clonally propagated, and the three events (3, 9, and 25) were effectively clones of the T0 events.

qPCR confirmed transgene expression in sorghum lines characterized in both the greenhouse and field. All three transgenic events showed high levels of *ZmRbcS* and *ZmRaf1* expression, with none in the WT (Fig. S3a-d). Immunoblots similarly showed high RAF1 protein expression in the transgenic lines relative to WT (Fig. 1b). Epitope tags showed expression of sorghum SS and RAF1 proteins in the sugarcane transformants (Fig. 1c).

**Fig. 1:**
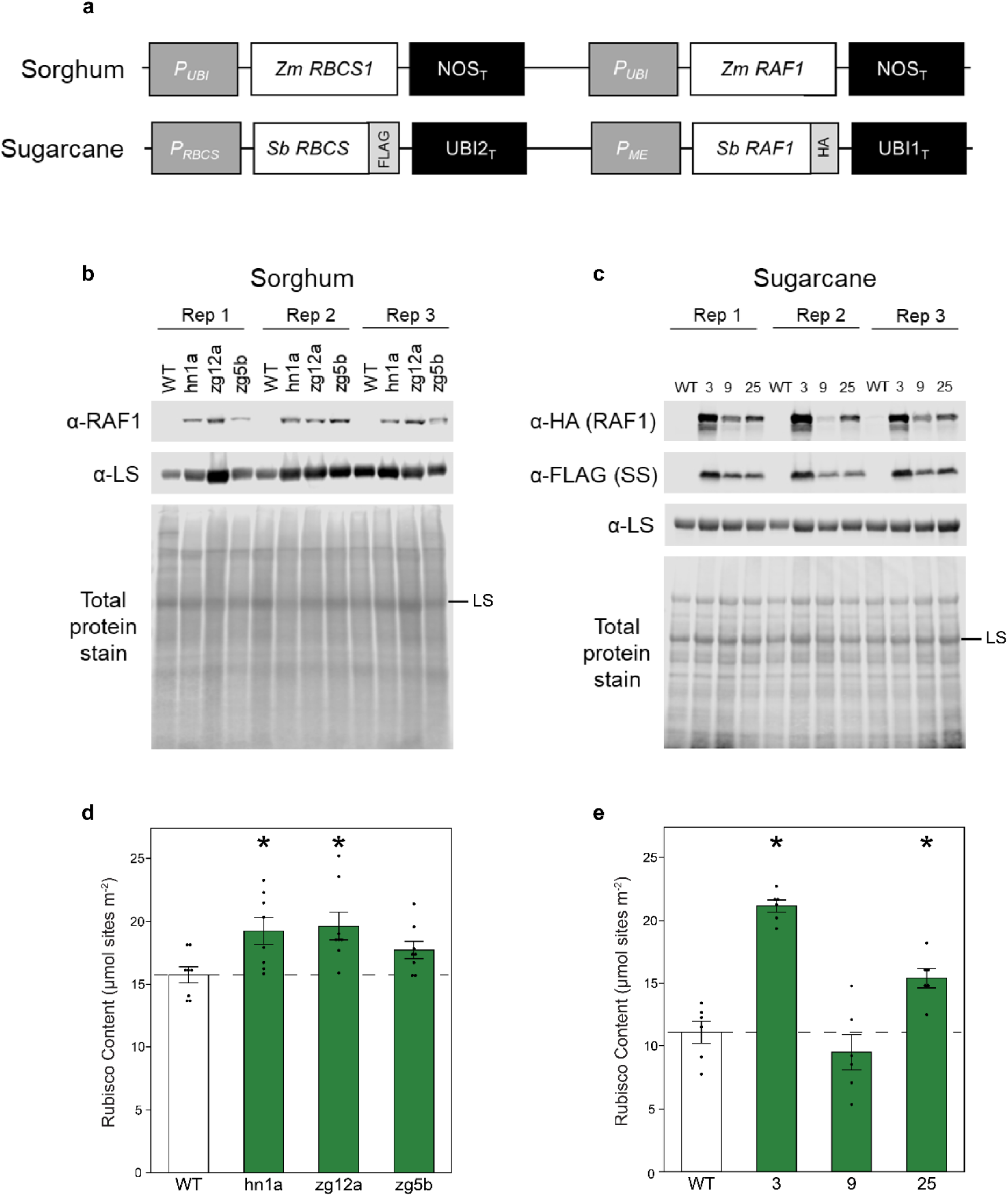
Sorghum and sugarcane transformation constructs, protein expression and analysis of Rubisco content. **(a)** Transgenes designed to upregulate Rubisco small subunit (*RbcS*) and Rubisco accumulation factor 1 (*Raf1*). The transgenes were stably transformed into sorghum (top) and sugarcane (bottom). The sugarcane construct contains FLAG and HA epitope tags. P, promoter; T, terminator, Sb, *Sorghum bicolor*; Zm, *Zea mays*; UBI, ubiquitin; NOS, nopaline synthase; ME, NADP-malic enzyme; HA, hemagglutinin. **(b)** Total soluble protein isolated on a leaf area basis from sorghum and **(c)** sugarcane, and analyzed by immunoblot. Three replicate plants (Rep) of each of the three transgenic events and of the wild-type (WT) control were probed with antibodies indicated to the left. -LS marks the Rubisco large subunit band. **(d)** Average Rubisco content per unit leaf area quantified by ^14^C-CABP binding in 6-week-old sorghum (n=8), and **(e)** 9-week-old sugarcane (n=6) grown in the greenhouse. Values are shown as the mean ± SEM. Asterisks indicate significant differences between WT and the transgenic line (*P < 0.05); one-way ANOVA, Dunnett’s post hoc test.

Rubisco content, measured by the ^14^C-CABP binding assay, was significantly increased by 13-25% in sorghum, and by 90% and 39% in sugarcane events 3 and 25 respectively (Fig. 1d,e). No significant increase was measured in event 9, even though immunoblots showed the presence of the recombinant SS and RAF1, although the latter at lower levels than in events 3 and 25 (Fig. 1c). There were no differences in total soluble protein between WT and transgenic plants in either sorghum or sugarcane (Table S3).

### Co-upregulation of RBCS and RAF1 is correlated with increased *V*_c,max_, CO_2_ assimilation, and photosynthetic light induction in sorghum

*A-C*_i_ responses showed that upregulated RBCS and RAF1 significantly increased assimilation rates above a *C*_i_ of ∼100 µmol mol ^−1^, but not below (Fig. 2a). Maximum *in vivo* Rubisco carboxylation rates (V_c,max_) at 25 °C, estimated from the *A-C*_i_ response, were significantly increased by 12% in the transgenic lines (Fig 2c), while maximum PEPC carboxylation rates (V_p,max_) though slightly higher than WT did not differ significantly (Table 1). *In vitro* V_c,max_ rates were increased by ∼40% in the transgenic lines (Fig. 2d). *A*_sat_ measured at the current atmospheric [CO_2_] also showed significant increases in the transgenic lines (Fig. 2b). *g*_sw_ was similarly significantly increased but this cannot account for increases in *A*_sat_ because no differences in *C*_i_ were observed between genotypes (Table 1). The effective quantum yield of PSII (Φ_PSII_), measured by modulated chlorophyll fluorescence, was increased by 12% in the transformed plants in parallel with CO_2_ assimilation (Table 1). Rubisco activation state, the fraction of Rubisco active sites that are catalytically active and therefore contributing to carboxylation, was not significantly different between lines (Table 1).

**Fig 2:**
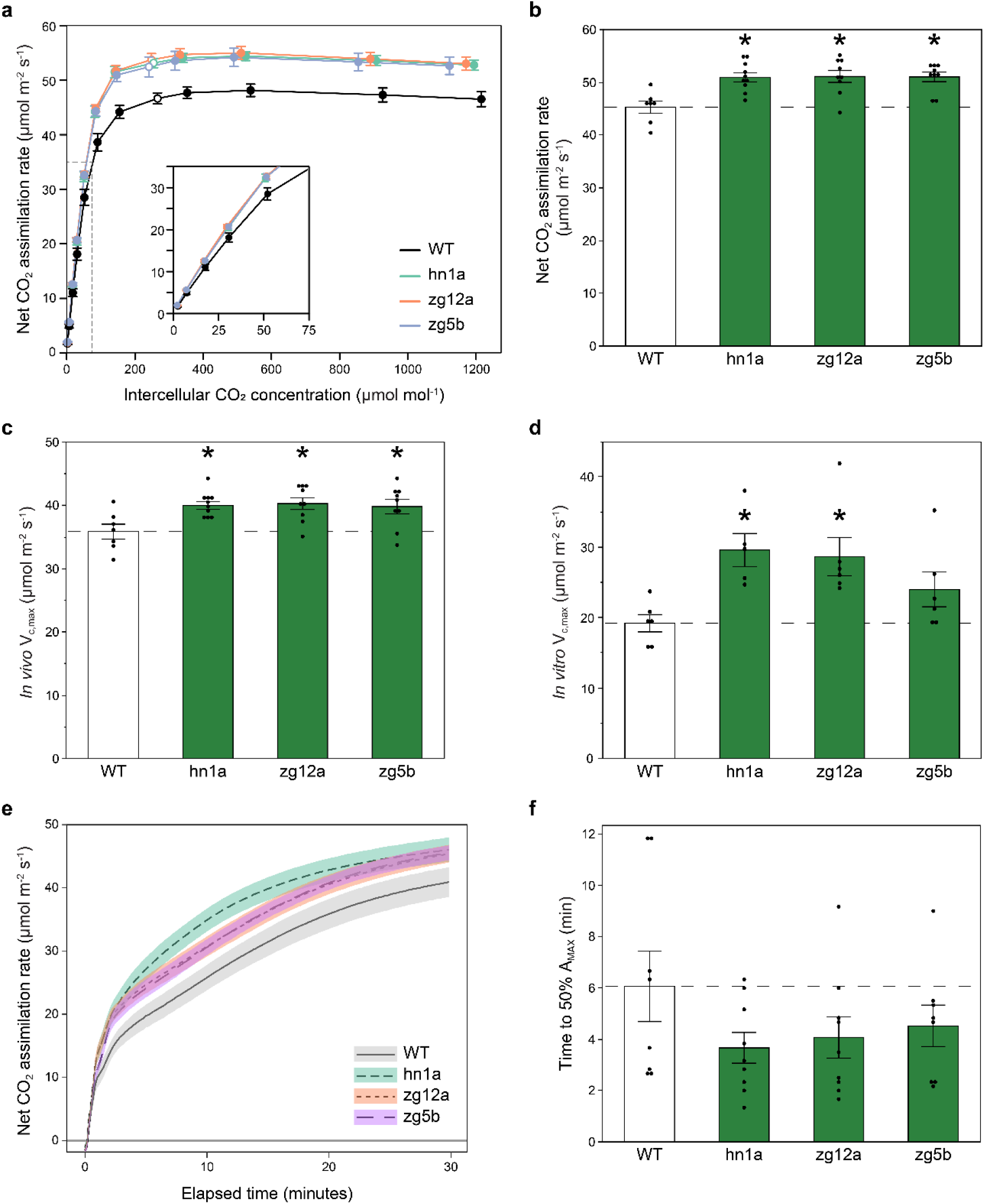
CO_2_ response curves and photosynthetic induction to light of greenhouse grown sorghum plants. **(a)** Response of light saturated CO_2_ assimilation rates (*A*_sat_) to intercellular [CO_2_] (C_i_). Measurements were made under the following conditions: light intensity of 1800 µmol m^−2^ s^−1^, leaf temperature of 28 °C, and VPD 1.5 kPa. Measurement cuvette [CO_2_] were varied from 10-1500 µmol mol^−1^ CO_2._ Open points represent the measured value closest to the operating *C*_i_ (i.e the *C*i where ambient CO_2_ was 420 µmol mol^−1^). **(b)** *A*_sat_ measured at 400 µmol mol^−1^ CO_2._ **(c)** Maximum *in vivo* Rubisco carboxylation rate (*V*_c,max_) at 25 °C estimated from response curves in panel **a** (n=7-10). **(d)** Maximum *in vitro* Rubisco carboxylation rate (*V*_c,max_) at 25 °C (n=5-6). **(e)** Average CO_2_ assimilation during the first 180 seconds of induction. **(f)** Time for *A* to reach 50% of *A*_max,_ where *A*_max_ is the average *A* from 25 to 30 minutes of illumination. Measurements were made on 5-week-old plants. Values are shown as the mean ± SEM (n=7-10). Asterisks indicate significant differences between WT and the transgenic line (*P < 0.05); one-way ANOVA, Dunnett’s post hoc test.

**Table 1.** Summary of biochemical and leaf gas exchange measurements from greenhouse grown sorghum and sugarcane. Values are shown as the mean ± SEM. Values significantly different from WT at *P* < 0.05 are in bold. One-way ANOVA, Dunnett’s post hoc test.

| Parameter | WT | RBCS-RAF1<br>Hn1a | RBCS-RAF1<br>Zg12a | RBCS-RAF1<br>Zg5b | Sample<br>number (N) |
| --- | --- | --- | --- | --- | --- |
| **Sorghum - Greenhouse** |  |  |  |  |  |
| Initial Rubisco Activity<br>($\mu\text{mol m}^{-2} \text{s}^{-1}$) | 13.48 ± 1.16 | **19.00 ± 1.45** | **22.28 ± 1.61** | 17.60 ± 1.20 | 5-6 |
| Rubisco Activation State<br>(%) | 71.41 ± 6.99 | 64.89 ± 4.47 | 79.21 ± 5.31 | 75.38 ± 5.79 | 5-6 |
| $\Phi\text{PSII}$ | 0.299 ±<br>0.002 | **0.336 ±**<br>**0.007** | **0.331 ±**<br>**0.009** | **0.338 ±**<br>**0.006** | 7-10 |
| $V_{p,\text{max}}$ ($\mu\text{mol m}^{-2} \text{s}^{-1}$) | 80.32 ± 7.38 | 88.33 ± 3.19 | 88.05 ± 3.59 | 89.99 ± 2.93 | 7-10 |
| $C_i$ ($\mu\text{mol mol}^{-1}$) | 153.53 ±<br>2.36 | 142.53 ±<br>2.18 | 144.62 ±<br>3.55 | 144.50 ±<br>4.95 | 7-10 |
| $g_{sw}$ ($\text{mol m}^{-2} \text{s}^{-1}$) | 0.411 ±<br>0.010 | **0.463 ±**<br>**0.009** | **0.472 ±**<br>**0.012** | **0.472 ±**<br>**0.014** | 7-10 |
| $\bar{A}_{180}$ ($\mu\text{mol m}^{-2} \text{s}^{-1}$) | 10.31 ± 1.17 | **14.25 ± 1.10** | **13.99 ± 0.90** | 12.83 ± 0.78 | 8-9 |
| Parameter | WT | RBCS-RAF1<br>3 | RBCS-RAF1<br>9 | RBCS-RAF1<br>25 | Sample<br>number (N) |
| **Sugarcane - Greenhouse** |  |  |  |  |  |
| Initial Rubisco Activity<br>($\mu\text{mol m}^{-2} \text{s}^{-1}$) | 15.89 ± 0.86 | **23.84 ± 2.84** | **23.74 ± 1.58** | 20.35 ± 1.81 | 6 |
| Rubisco Activation State<br>(%) | 82.30 ± 4.38 | 81.86 ± 4.54 | 87.22 ± 3.10 | 79.40 ± 7.51 | 6 |
| $\Phi\text{PSII}$ | 0.297 ±<br>0.012 | 0.319 ±<br>0.008 | 0.300 ±<br>0.009 | 0.326 ±<br>0.009 | 10 |
| $V_{p,\text{max}}$ ($\mu\text{mol m}^{-2} \text{s}^{-1}$) | 68.82 ± 4.55 | 85.70 ± 6.97 | 68.99 ± 3.02 | 79.93 ± 4.76 | 9-10 |
| $C_i$ ($\mu\text{mol mol}^{-1}$) | 116.48 ±<br>8.61 | 113.76 ±<br>6.29 | 121.17 ±<br>4.47 | 118.55 ±<br>3.27 | 9-10 |
| $g_{sw}$ ($\text{mol m}^{-2} \text{s}^{-1}$) | 0.252 ±<br>0.012 | 0.282 ±<br>0.006 | 0.282 ±<br>0.007 | **0.305 ±**<br>**0.015** | 9-10 |
| $\bar{A}_{300}$ ($\mu\text{mol m}^{-2} \text{s}^{-1}$) | 11.05 ± 2.26 | 17.78 ± 2.08 | 5.62 ± 1.95 | 15.38 ± 1.77 | 7-8 |
| Time to 50% $A_{\text{MAX}}$ (min) | 3.36 ± 0.38 | 3.05 ± 0.27 | **9.27 ± 1.62** | 3.35 ± 0.26 | 6-8 |

To identify if increased Rubisco would be beneficial to the activation of photosynthesis on shade to sun transition, induction of *A* over the first 30 min of transfer to high-light (PPFD = 1800 µmol m^−2^ s^−1^) was measured (Fig. 2e). CO_2_ assimilation in the transgenic lines was significantly higher than WT after 3 minutes of illumination following dark acclimation (Table 1) and remained higher for the remaining 27 minutes of measurement (Fig. 2e). Induction curves, normalized to the average *A* value over the last 5 minutes of illumination (*A*_max_) were used to assess the speed of photosynthetic activation (Fig. S4a). On average, the transgenics took 4 minutes to reach 50% maximum CO_2_ assimilation rates (*A*_max_), compared to 6 minutes for WT (Fig. 2f). The transgenics were significantly taller and had more leaves than WT in the greenhouse (Fig. S5). Seed from these plants was collected for the field trial.

### RBCS-RAF1 transgenic sorghum plants have increased photosynthesis and biomass under field conditions

A field trial in summer 2023 evaluated whether gains observed in the greenhouse could be reproduced in a replicated field trial of 4-row plots. A randomized complete block design was used with 6 replicate plots for each of the three RBCS-RAF1 transgenic events and WT (Fig. 4e; Fig. S6a). *A-C*_i_ responses and light induction curves measured during the vegetative growth stage were consistent with the greenhouse results. *V*_c,max_, was again increased, with no significant increase in the initial slope of the response, and light induction of *A* was faster (Fig 3a,b). These increases were also observed at boot stage but were absent during grain-filling (Fig. 3c,d). Bundle sheath leakiness (*Ф)*, the rate that CO_2_ not fixed by Rubisco leaks from bundle sheath back to mesophyll, was measured by combining gas exchange with carbon isotope discrimination during the vegetative growth stage. In concert with increased *V*_c,max_, *Ф* was decreased by about 18% across all three transgenic lines, however these decreases were not statistically significant (Fig. S7).

**Fig. 3:**
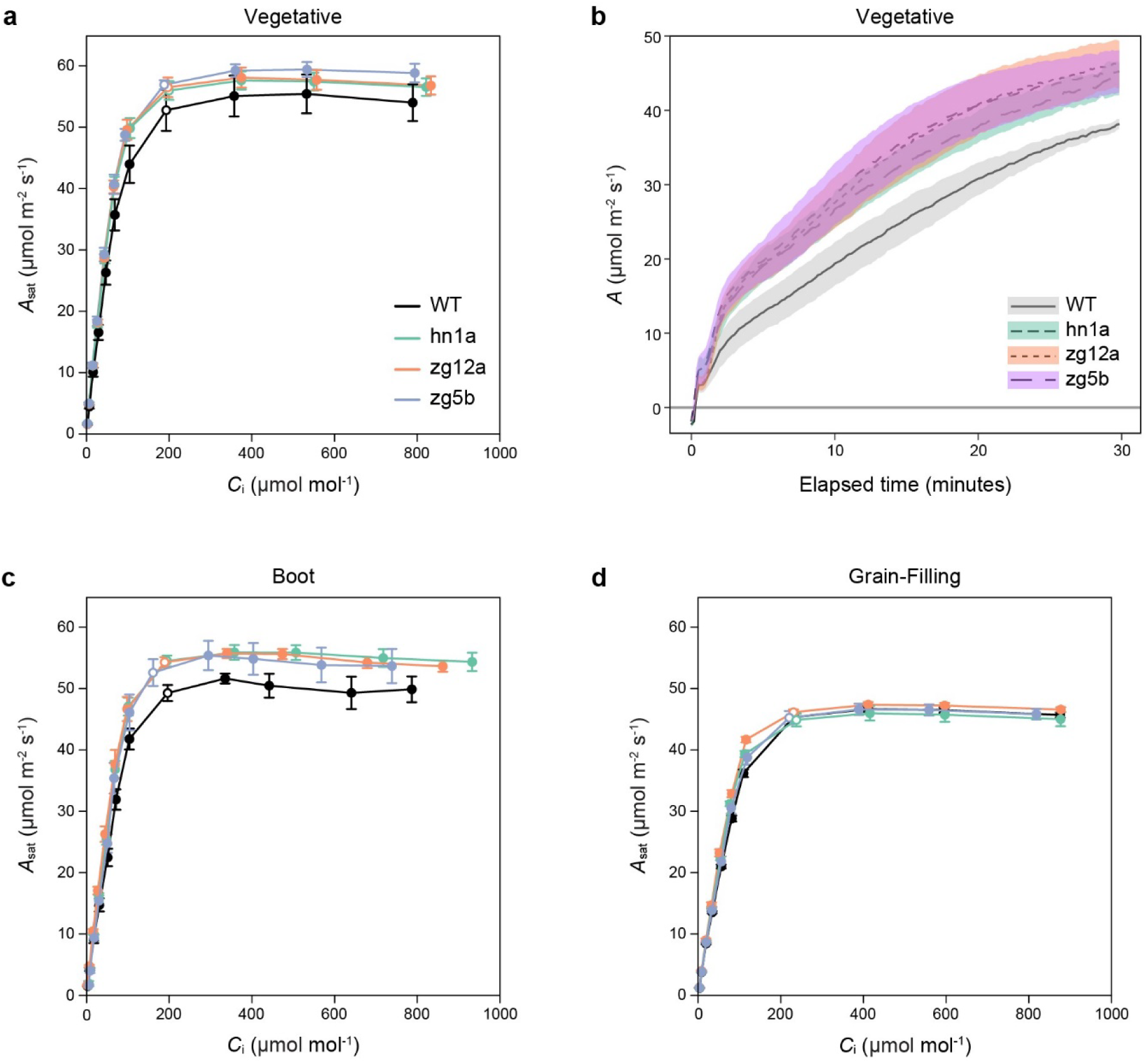
CO_2_ response curves and photosynthetic induction to light of field grown sorghum plants. **(a)** Response of light-saturated CO_2_ assimilation rates (*A*_sat_) to intercellular [CO_2_] (*C*_i_) and **(b)** induction of *A* during the first 30 min of illumination at PPFD 1800 µmol m^−2^ s^−1^ of sorghum plants in the vegetative growth stage. **(c)** *A*_sat_ *-C_i_* response of sorghum plants in the boot and **(d)** grain-filling growth stages. Open symbols on each curve show the measured value closest to the operating C_i_ (where ambient CO_2_ is 420 µmol mol^−1^). Values are shown as the mean ± SEM (n=4-6).

**Fig. 4:**
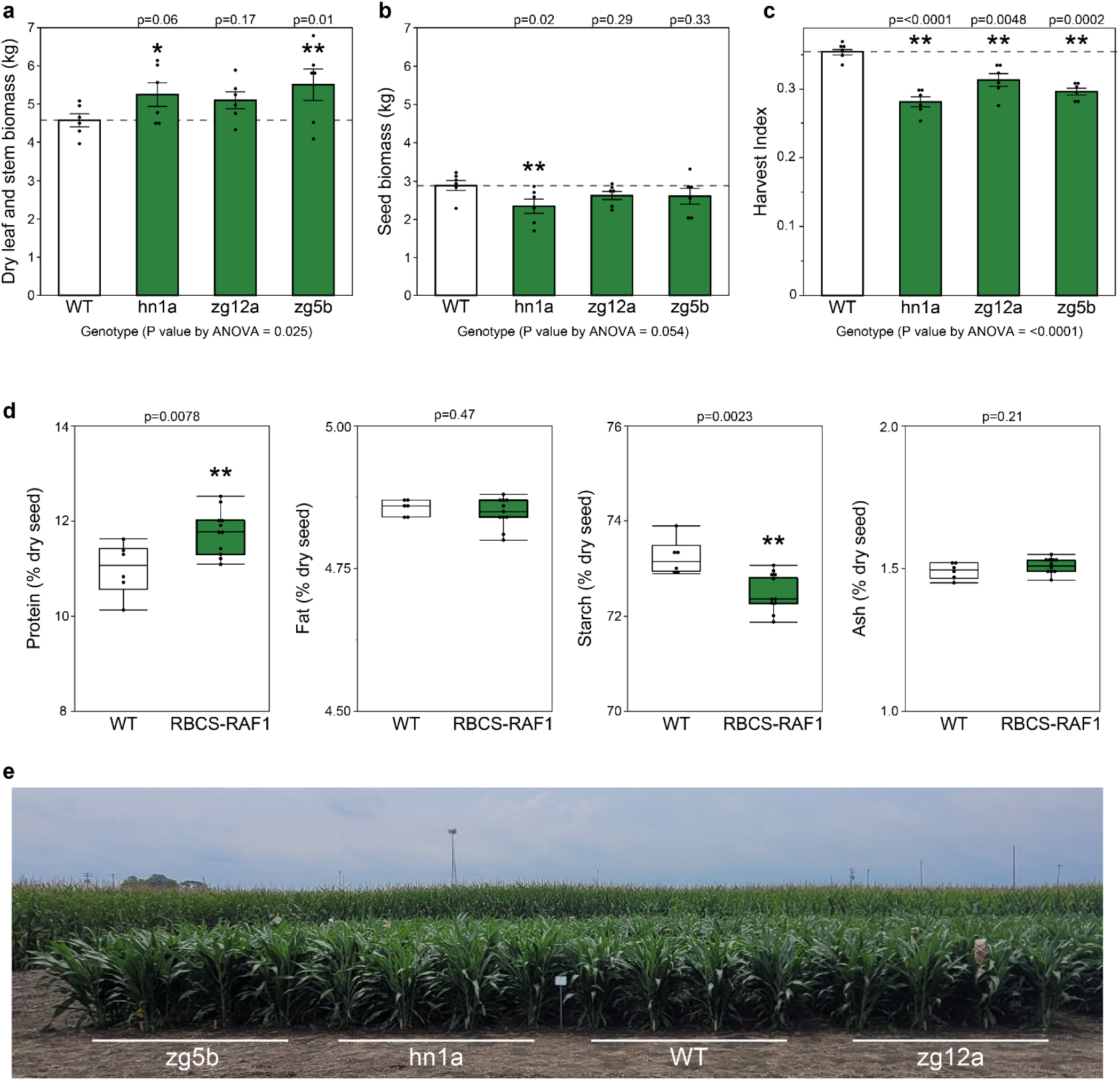
Plant growth traits in field grown sorghum plants. **(a**) Average dry leaf and stem biomass per plot and **(b)** average seed biomass per plot from field grown sorghum plants. **(c)** Harvest index (seed mass/total shoot mass) per plot. Values are shown as the mean ± SEM (n=6 plots). Asterisks indicate significant differences between WT and the transgenic line (**P < 0.05, *P < 0.1); one-way ANOVA, Dunnett’s post hoc test. P values above each bar from Dunnett’s post hoc test. **(d)** Percent protein, fat, starch and ash content of whole dry sorghum seeds (WT n= 6 plots, RBCS-RAF1 n=11 plots). **P < 0.05; pooled t-test. **(e)** Side view of 4 of the 24 four-row plots of the four sorghum genotypes tested at boot stage in the field in Urbana, Illinois summer 2023 (see Fig. S4a for full experimental design).

The field grown transgenic sorghum plants were also more vigorous than the WT, as evidenced by a significant 15.5% increase in vegetative dry biomass at harvest (Fig. 4a). However, there was no increase in seed biomass, which corresponded to a significant decrease in harvest index (seed mass/total plant mass) compared to the WT (Fig. 4b-c). Seed composition was analyzed using NIR and showed increased protein and decreased starch content in the transgenic lines, with no change in fat or ash content (Fig. 4d). Number of panicles per plot, 100 seed weight and average seed weight per panicle did not show any clear trends, however panicle emergence was delayed by 3 to 6 days in the transgenic plants (Table S2). Leaf mass, chlorophyll content, N and C contents, on a leaf area basis, did not differ between the WT and transgenic plants (Table S3).

### Co-upregulation of RBCS and RAF1 in sugarcane increases photosynthetic efficiency and plant productivity

Measurements were made on 8-week-old greenhouse grown sugarcane plants. As in sorghum, *A-C*_i_ responses showed significant increases in *in vivo* Rubisco activity (*V*_c,max_) with no significant change in the initial slope (*V*_p,max_, Fig. 5a,d; Table 1). Similarly, *A*_sat_ and *in vitro V*_c,max_ were significantly increased at ambient atmospheric [CO_2_] in the transgenics (Fig. 5c,e). There were no significant differences in Rubisco activation state, *g*_sw_ or *C*_i_: however, there was a marginal increase in Φ_PSII_ in events 3 and 25 (Table 1). Assimilation of CO_2_ was greater over the 30 minute light induction, in events 3 and 25, but not event 9 potentially due to a pleotropic effect related to the location of the transgene insertion into the genome (Fig. 5b, Fig. S8).

**Fig 5:**
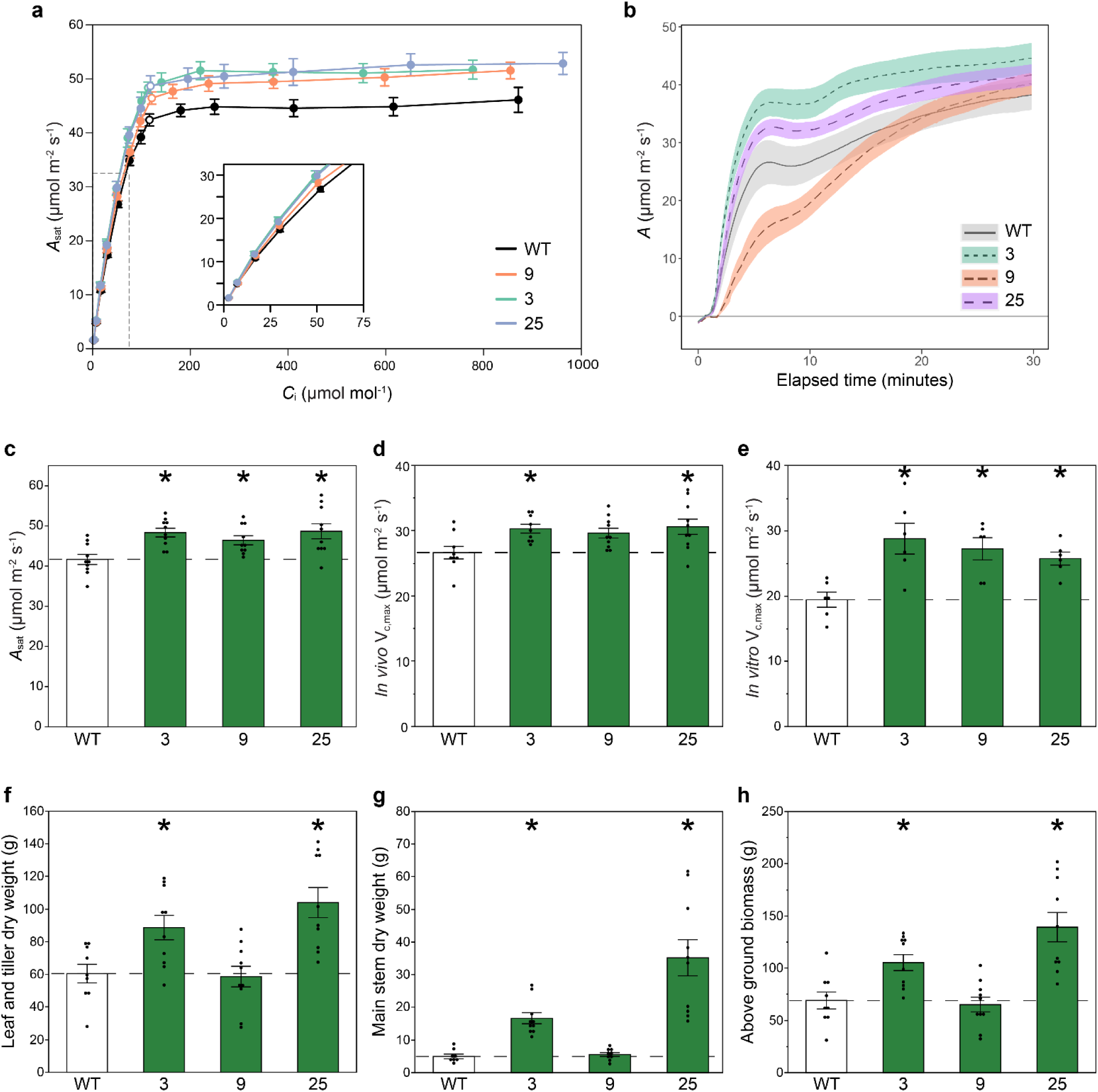
CO_2_ response curves and plant growth traits of greenhouse grown sugarcane plants. **(a)** Response of light-saturated CO_2_ assimilation rates (*A*_sat_) to intercellular [CO_2_] (*C*_i_). Measurements were at light intensity of 1800 µmol m^−2^ s^−1^, leaf temperature 30 °C, and 65% humidity. Measurement cuvette [CO_2_ ] was varied from 10-1500 µmol mol^−1^. Open symbols show the measured value closest to the operating C_i_ (where ambient [CO_2_] was 420 µmol mol^−1^). **(b)** Induction of *A* during the first 30 min of illumination at PPFD 1800 µmol m^−2^ s^−1^. **(c)** *A*_sat_ at 450 µmol mol^−1^ CO_2_ **from panel a. (d)** Maximum Rubisco carboxylation rate *in vivo* (*V*_c,max_) at 25 °C estimated from response curves in panel **a** (n=8-10). **(e)** Maximum *in vitro* Rubisco carboxylation rate (*V*_c,max_) at 25 °C (n=6). **(f)** Leaf and tiller, **(g)** main stem, and **(h)** above ground total dry biomass of greenhouse grown sugarcane plants. Values are shown as the mean ± SEM (n=8-10). Asterisks indicate significant differences between WT and the transgenic lines (*P < 0.05); one-way ANOVA, Dunnett’s post hoc test.

Transgenic events 3 and 25 showed significant increases in main stem, and tiller and leaf dry mass, amounting to total shoot dry mass increases of 37% and 81% respectively, but with no increase in event 9 (Fig. 5f-h). Significant increases in plant height, stem width and leaf number were also observed in events 3 and 25 compared to WT (Fig. S9). Leaf chlorophyll and mass per unit area did not differ between genotypes (Table S3).

## Discussion

Theoretical and experimental analyses have long suggested Rubisco to be the major limitation of light-saturated photosynthesis in both C_3_ and C_4_ species (7, 18–20, 28). Consistent with this hypothesis, transgenic upregulation of Rubisco content significantly increased Rubisco carboxylation rates (*V*_cmax_) and light-saturated leaf CO_2_ assimilation rates (*A*_sat_) in both sorghum and sugarcane (Fig. 1 d-e, 2a-d, 3a-c, and 5a, c-e). Although Rubisco or electron transport can limit assimilation in the plateau of a C_4_ *A*-*C_i_* curve (7, 29), the agreement between *in vivo* and *in vitro V_cmax_* estimates is consistent with primarily Rubisco-limited assimilation at high light (Fig. 2a,c-d and 5 a, d-e). The speed of light induction of CO_2_ assimilation was also increased (Fig. 2e-f and 3b). This has critical importance to increasing photosynthetic efficiency under the fluctuating light conditions of crop canopies in the field (30, 31). Also consistent with limitation by Rubisco, bundle sheath leakiness was decreased, although not significantly (Fig. S7). Finally, and in concert with these increases in photosynthetic efficiency, a field trial of sorghum and a greenhouse trial of sugarcane showed significant increases in productivity relative to the wild-type from which the transgenics were derived (Figs. 4 and 5).

Rubisco is, by far, the most abundant single protein in leaves, constituting up to 50% of soluble protein in C_3_ plants and 20-30% in C_4_ (32). These high amounts are attributed to the need to offset the exceptionally low catalytic rate of carboxylation (*k*_cat_) of Rubisco relative to other enzymes. Its low *k*_cat_ is in turn considered in part a consequence of high specificity for CO_2_ relative to the alternative substrate of O_2_ (33). Sage & McKown (34) noted the much lower available tissue volume that could house Rubisco in C_4_ leaves, where it is functionally confined to the bundle sheath, suggesting this as a constraint. This is less than half the volume available in C_3_ leaves. However, 3D microscopic analysis of the bundle sheath chloroplast stromal volume in *Miscanthus* x *giganteus* suggested capacity for a doubling of actual Rubisco content (35). *Miscanthus* is closely related to sorghum and sugarcane, and within the same subtribe, Saccharinae (36). This observation is consistent with the observed increases in Rubisco content of 25% in sorghum and up to 90% in sugarcane (Fig. 1d-e). The much higher level in sugarcane may reflect that this was achieved by biolistic transformation, which could have resulted in multiple insertions of the transgenes, while the three sorghum events were single insertion events.

Given that increased Rubisco production is possible and beneficial, why have natural selection and breeding not achieved this? One explanation is that Rubisco might only have become a limitation very recently, due to rising atmospheric [CO_2_]. On March 16, 2024, atmospheric [CO_2_] was recorded at 428 μmol mol^−1^ (21), which is almost double the average atmospheric [CO_2_] of the past 40,000 years (22, 37). Theory suggests that under the conditions in which our crop ancestors evolved, Rubisco would not be a limitation and therefore selection pressure would have been lacking, with the result that there may be little genetic variation for natural or breeder selection as [CO_2_] rises. Indeed given the large nitrogen cost of Rubisco, historically, selection pressure was likely against any increase. It should be noted that half of the increase in [CO_2_] has occurred in the last 50 years, with [CO_2_] rising from 323 μmol mol^−1^ to 428 μmol mol^−1^between 1974 and 2024 (21). By reference to the open symbols in Fig. 3, it is seen that at today’s atmospheric [CO_2_] *A*_sat_ in the field-grown sorghum is controlled by the plateau of the *A*-*C*_i_ response, i.e. predominantly by Rubisco activity. However, if it is assumed that *C*_i_/*C*_a_ has been conserved, then at the atmospheric [CO_2_] in 1974, *C*_i_ would equal 129 μmol mol^−1^. By reference to Fig. 3, *A*_sat_ would then be mostly controlled by the initial slope of the *A*-*C*_i_ response, i.e. PEP carboxylase activity (*V*_p,max_), and insensitive to increase in Rubisco activity. The period in which Rubisco has become limiting is therefore very short in terms of reproductive cycles. As atmospheric [CO_2_] continues to rise, Rubisco limitations will increase, and elevation of Rubisco will be an important means to contribute toward future-proofing and advancing C_4_ crop productivity. An increase in *A*_sat_ was achieved here by only upregulating expression of *RbcS* and *Raf1,* which infers three factors important for future manipulations. First, for Rubisco to be increased, both the large (LS) and small (SS) subunits must be increased. That only upregulation of *RbcS* was needed, is consistent with the observations that in tobacco *rbcL* is subject to assembly-dependent translational regulation, and so controlled by the supply of SS (38) or in rice the supply of SS upregulates *rbcL* transcript levels (39). Further studies are required to determine which mechanism is occurring in sorghum and sugarcane. Second, although all three transgenic sugarcane events showed expression of the *Sb* SS protein, in event 9 this did not translate into improved productivity (Fig. 1c-e and 5f-h). Expression of *Sb* RAF1 was lowest in this line, suggesting the importance of effective expression of RAF1 with SS (Fig.1c). Third, there must either be 1) over-capacity in the C_4_ cycle, 2) the C_4_ cycle increases capacity in concert with increased Rubisco activity, or 3) both occur. The fact that bundle sheath leakiness of CO_2_ decreased and *V*_p,max_ slightly increased with increased Rubisco is consistent with hypothesis 3 (24).

Similar to results reported in maize (26), increases in Rubisco content corresponded to increases in Rubisco enzyme activity, resulting in higher photosynthetic rates and larger biomass. Activation of Rubisco requires the chaperone protein Rubisco activase (RCA). In contrast to previous studies in maize and rice (26, 40), no decrease in the fraction of active Rubisco was observed in the transgenic plants, indicating that sorghum and sugarcane must have had sufficient RCA to support the activation of all their additional Rubisco (Table 1). In these previous studies, increased Rubisco content was achieved by overexpressing endogenous *RbcS* with or without *Raf1*. Here, *RbcS* and *Raf1* genes from maize and sorghum were expressed in sorghum and sugarcane respectively, so the transgenic plants are assembling some chimeric Rubiscos comprised of the foreign transgenic SS and endogenous LS. Maize, sorghum and sugarcane have been reported to have very similar *k_cat_* (41, 42), implying that the increases in Rubisco activity reported here are solely due to increased Rubisco active sites and not changes in Rubisco kinetic properties. In addition, the FLAG tag fused to *RbcS* in the sugarcane construct did not appear to interfere with Rubisco activity (Fig. 5e).

In dense modern crop canopies, only the uppermost leaves might experience light saturated conditions. Most leaves will experience frequent light fluctuations, or sunflecking, due to the passage of clouds, shadowing by overlying leaves as solar angle changes and wind driven movements (30, 31). The speed with which photosynthetic efficiency can adjust to light fluctuations has been shown to have a profound effect on whole crop daily CO_2_ assimilation and productivity (43, 44). A key factor here is the speed of induction of *A* on transfer of a leaf from shade to sun. Modeling and experimental studies have shown the speed at which Rubisco activity increases to be critical to induction in both C_3_ and C_4_ crops (23, 45, 46). Here, increased Rubisco content coincided with more total CO_2_ assimilation over the 30 minutes of induction in all three sorghum events and two of the sugarcane events (Fig. 2e, 3b, 5b). Speed of induction, measured as time to 50% *A*_max_, was also increased in the transgenic sorghum lines but not in sugarcane (Fig. 2f and Table 1). This is consistent with computational results from Wang *et al.* (2021), which suggest that Rubisco activation is the main limiting factor in the speed of photosynthetic induction in sorghum, while the speed of stomatal opening and the concentration of the PPDK regulatory protein exert more control in sugarcane.

In the greenhouse, significant increases in growth of both sorghum and sugarcane were observed at an individual plant level (Fig. 5f-h; Fig. S5). Importantly, and consistent with increased CO_2_ assimilation rates during vegetative and boot stages, leaf and stem biomass was significantly increased by an average of 15.5% in the RBCS-RAF1 sorghum field plots relative to the WT controls, a desired characteristic for C_4_ biomass crops (Fig. 4a). However, grain yield per plot did not increase, due to a significant decrease in harvest index (Fig.4c). Although sorghum cultivar RTx430 has become the standard for transformation and experimental analyses, it is a 40-year old inbred line (47–49) and might be sink-limited. There is similar evidence for sink limitations in a 25-year old hybrid cultivar; when sorghum was grown under well-watered conditions under open-air CO_2_ enrichment using FACE technology, photosynthetic capacity and photosynthetic rates were increased (50–52). Yet, as in our experiment, stover increased while harvest index declined by 7% (53). Sink limitation at grain filling results in accumulation of non-structural carbohydrates in the source tissues, causing down-regulation of photosynthetic genes, notably *RbcS* (54). The loss of *in vivo* Rubisco activity (*V*_c,max_) during grain filling evident in Fig. 3d is consistent with sink limitation. Alternatively, grain yields may not have increased due to alternative factors such as delayed panicle emergence or decreased transgene expression. It is also important to note that Rubisco levels increased without an associated change in total soluble protein, LMA or leaf N content (Fig. 1 and Table S3). This suggests that more leaf N Is invested into Rubisco in the transgenic plants and therefore less N allocated to other leaf proteins. Although there was no negative impact on photosynthesis, altered N use may affect other processes including grain filling. Testing whether greater grain yields can be obtained will require the production of hybrids and/or transformation of current elite cultivars. Upregulation of Rubisco has been achieved here, as a test of the concept that more Rubisco would translate to more efficient photosynthesis and productivity in today’s atmosphere, by addition of extra copies of *RbcS* and *Raf1.* Alternatively, a future approach for upregulating Rubisco could be using DNA editing to mutate repressive elements in the promoter regions of the native *RbcS* and *Raf1* genes.

Despite significant breeding efforts, the average yield of sorghum has not changed over 30 years, nor has the yield of sugarcane in the USA in the last 20 years (Fig. S10a,b). This might reflect increasingly adverse weather conditions. However, even in the USA where it might be assumed that favorable agronomy would allow the yield potential to be realized, there is little evidence of yield improvement since 1991, even in the most favorable years (Fig. S10a). Therefore, if the photosynthetic and productivity improvements found here by upregulating Rubisco translate into parallel yield improvements in elite cultivars and hybrids, this could be very useful for integration into breeding programs. This successful test-of-concept suggests multi-location and/or multi-year replicated yield trials of Rubisco upregulation in elite material as the next proving step. Given the great value and global significance of this key economic clade of crops both for food and biomass, the results here indicate an important opportunity to achieve increased productivity under rising atmospheric [CO_2_].

## Methods

### Sorghum plasmid assembly and transformation

A 3.6 kb fragment containing the *Raf1* expression cassette regulated by the maize ubiquitin promoter was isolated from the plasmid pPTN1063 (26) by digesting the vector with *Kpn*I and *Hind*III. The fragment was then blunted with T4 DNA polymerase. Plasmid pPTN438 (38) (containing the *RbcS* expression cassette) was linearized via *Kpn*I digestion and blunted with T4 DNA polymerase. The pPTN1063 fragment and linearized pPTN438 plasmid were then ligated resulting in the final binary vector pPTN1589. Sorghum inbred RTx430 was transformed with pPTN1589 using agrobacterium as previously described (55).

### Sugarcane plasmid assembly

Recombinant DNA constructs were assembled with Golden Gate cloning to highly express Rubisco small subunit (*RbcS*) and Rubisco accumulation factor 1 (*Raf1*) from *Sorghum bicolor* in sugarcane. The open reading frame (ORF) of *SbRbcS* (Sobic.005G042000) and ORF of *SbRaf1* (Sobic.006G269400) were synthesized with 1X FLAG tag and 6X HA tag, respectively (GenScript, Piscataway, NJ). *SbRbcS*-FLAG was subcloned under transcriptional control of the maize RBCS promoter + 5’UTR and the *Panicum virgatum* UBI2 (PvUBI2) terminator. The synthesized *SbRaf1*-HA was subcloned under transcriptional control of the maize NADP-ME promoter and *Panicum virgatum* UBI1 (PvUBI1) terminator. The selectable marker gene *npt*II was subcloned under transcriptional control of the ubiquitin promoter from *Zea mays* and SbHSP16.9 terminator from *Sorghum bicolor*. The three expression cassettes were subcloned into a L2 construct downstream of an RB7 matrix attachment region (MAR) and upstream of TM2 MAR from *Nicotiana tabacum* serving as insulators and confirmed by Sanger sequencing. The minimal expression construct flanked by MARs was excised from the plasmid backbone using the *I-Sce*1 restriction enzyme and gel purified prior to biolistic transformation into sugarcane.

### Sorghum transgene copy number analysis

Transgene copy number was measured using digital droplet PCR (ddPCR) following the protocols from Głowacka et al. (56), with slight modifications. See Table S1 for primers used (57). Sorghum genomic DNA was digested overnight with HindIII. A 20 μL ddPCR reaction mix contained 20 ng of the digested DNA, 10 μL QX200 ddPCR EvaGreen Supermix (186-4034, BioRad, Hercules, CA) and primers at a final concentration of 100 nM. Droplets were generated using a QX200 droplet generator (186-4002, Bio-Rad) by combining 20 μL of the reaction mix with 70 μL of EvaGreen droplet generation oil (186-4005, Bio-Rad). The PCR reaction was conducted in a deep-well thermal cycler (C1000 Touch, 185-1196, Bio-Rad) with the following conditions: 95 °C for 5 minutes, 40 cycles of 95 °C for 30 seconds and 59 °C for 1 minute, then 4 °C for 5 minutes and 90 °C for 5 minutes, with a ramp rate of 2.5 °C/second at each step. Immediately after the PCR, measurements were taken using a QX200 droplet reader (186-4003, Bio-Rad).

### Generation and PCR confirmation of transgenic sugarcane plants

The linearized, minimal construct containing *SbRbcS*, *SbRaf1* and NPTII cassettes flanked by matrix attachment regions was introduced into sugarcane cv. CPCL02-0926 by biolistic gene transfer as described previously (58). Briefly, cross-sections of sugarcane leaf whorls above the shoot apical meristem were cultured on a modified Murashige & Skoog medium (MS) with B5 vitamins (PhytoTech Labs, KS, USA) supplemented with 3 mg/L of 2,4-Dichloro phenoxy acetic acid (PhytoTech Labs) and maintained at 28 ℃ in the dark for induction of embryogenic callus. For biolistic gene transfer, the minimal expression cassette was coated onto gold particles and introduced into embryogenic calli using biolistic PDS-1000/He delivery system (Bio-Rad) as described previously (59). Following 4 weeks of selection of transformed calli on the medium containing the antibiotic Geneticin®, calli were sub-cultured onto regeneration media (MS media with B5 vitamins, supplemented with 1.86 mg/L of α-Naphthaleneacetic acid/NAA (PhytoTech Labs) and 0.09 mg/L of 6-Benzylaminopurine/BAP (PhytoTech Labs)) and maintained at 28 ℃ incubator with 16/8 hr light (100 µmol m^−2^ s^−1^) and dark cycle. Regenerated shoots were transferred to modified MS basal medium (PhytoTech Labs) for shoot elongation and rooting. Rooted plantlets were transferred to soil and acclimatized in a temperature-controlled growth chamber maintained at 28 ℃ day and 22 ℃ night temperature with 16/8 hr light (400 µmol m^−2^ s^−1^) and dark cycle for growth and further analysis.

For the PCR confirmation of transgenic events, DNA was isolated from 100 mg of leaf tissues using CTAB method with modifications (60). PCR reactions were performed on a thermocycler (Applied Biosystems, Foster City, CA) using standard Taq buffer, dNTPs and hot start Taq polymerase (New England Biolabs Inc, Ipswich, MA). See Table S1 for primers used for PCR amplification. Amplicons were resolved with electrophoresis on a 1.2% agarose gel and visualized under UV transilluminator (Bio-Rad).

### Plant growth – greenhouse conditions

T2 homozygous *Sorghum bicolor* (inbred RTx430) seeds from 3 independent single insertion events and non-transformed WT seeds from the same harvest date were sowed into plastic potting trays 6 cm x 6 cm x 5.4 cm deep, filled with BM7 growing medium (BM7 All-Purpose Bark Mix, Berger). After about 14 days seedlings were transplanted into 9.5L plastic pots filled with BM7 growing medium supplemented with 44 cm^3^ of 13-13-13 (N-P-K) granulated slow-release fertilizer (Osmocote, ICL-Growing Solutions). Sugarcane (*Saccharum officinarum* cv. CPCL02-0926) plantlets clonally propagated from three independent T0 transformation events and non-transformed WT plants were also grown in the greenhouse. Single node segments from mature stalks were transplanted into 9.5L plastic pots filled with BM7 growing medium supplemented with 44 cm^3^ of 13-13-13 (N-P-K) granulated slow-release fertilizer. Both sorghum and sugarcane were grown under natural illumination (Figure S11). Lights turned on from 6:30am to 9:30pm (15 hour days), providing an additional 350 to 450 µmol m^−2^s^−1^ of LED light, and turned off when outside light intensity surpassed 350 watts/m^2^ (∼800 µmol m^−2^s^−1^). The greenhouse temperature was kept between 25-28°C in the day and 24-26 °C at night. Both sorghum and sugarcane plants were fertilized with a rotation of 20-10-20 & 15-5-15 liquid fertilizer (JR Peters INC) at 300 ppm N twice a week. Once a month a MgSO_4_ (100 ppm) and iron (10 ppm) drench was applied (Sprint 138, BASF Corporation).

Sugarcane plants were harvested after 2 months of growth. Immediately prior to harvesting, leaf number (fully expanded), tiller number, plant height and stem width were measured. Stem width was measured at the widest point approximately 5 cm above the soil using calipers. Height was measured from the soil to the top of the youngest fully expanded leaf (leaf collar present). Leaf mass per area (LMA) was measured from 3.15 cm^2^ of leaf discs, which were dried until constant weight and weight recorded. Chlorophyll content was measured using a SPAD meter (502, Spectrum Technologies). Harvested material was partitioned into main stem biomass and leaf plus tiller biomass and dried at 60°C to steady state weight. Above ground biomass was calculated as the sum of stem, leaf and tiller biomass.

### Plant growth – field conditions

Sorghum seed obtained from the plants in the greenhouse experiment were harvested and used for a field experiment. A randomized complete block design with six blocks was used. Each block contained 16 rows in a North-South direction, made up of four row plots from each of the transgenic lines and WT control (Fig. S6a). Prior to planting the ground was worked with a field cultivator (April 27 2023), and no fertilizer was applied to the soil. Seeds were sowed using a precision planter at a density of 8.6 m^−2^ (Almaco) on June 2^nd^ 2023 on the Crop Sciences Farm of the University of Illinois SoyFACE Experimental Station (40°02’36.0“N 88°13’48.5“W). Weather data was collected with an automatic weather station on the Farm (Fig. S6b-d). July 5^th^ Warrior II with Zeon technology (Syngenta) was applied at a moderate rate of 0.07 dm^3^/ha for the control of flea beetles. August 2^nd^ Mustang Maxx (FMC Corporation) was applied at moderate rate of 0.1 dm^3^/ha for the control of aphids.

To avoid any release of pollen, paper bags enclosed the panicles, sealed at the peduncle, prior to anthesis, and remaining until harvest (recorded as panicle emergence date). Plants were harvested by hand on October 4^th^ from the middle two rows of each plot by cutting stems at ground level. Two plants from either end of each row were discarded to avoid edge effects. Harvested material was partitioned into stem/leaves and panicles with seed. Stem and leaves were dried to a constant weight in custom made drying ovens at 60°C and panicles with seed were dried at 30°C. Dry weights per plot were obtained for both, then seed was collected using a threshing machine (BT 14-E Belt Thresher, Almaco). The machine was cleaned between each sample to avoid contamination. Total seed weight per plot, and 100 seed weights were then determined. Above ground biomass was calculated as the sum of stem, leaf, panicle and seed biomass. Harvest index was calculated as seed biomass divided by total above ground biomass. Seed weight per panicle was estimated by dividing seed weight per plot by panicle number per plot. Seed protein, starch, fat and ash content were estimated on a percent basis using a near-infrared reflectance (NIR) analyzer (DA 7250, Perten Instruments). Seeds were placed into the sample dish and analyzed using a built in program for whole-seed evaluation (“Sorghum-Grain”). Chlorophyll content was measured using a SPAD meter (502, Spectrum Technologies) on August 30^th^. LMA was measured from from 6.06 cm^2^ of leaf tissue collected August 31^st^

### Transcript and protein expression

Prior to RNA and protein extraction tissue was disrupted and homogenized as described previously (61). RNA was extracted from 1.9 cm^2^ of leaf tissue from the youngest fully expanded leaf of greenhouse and field-grown sorghum, using the NucleoSpin RNA Plant Kit (Macherey-Nagel). 700 µL of the lysis buffer was used. RNA quantity and quality was assessed using a NanoDrop (Thermo Fisher Scientific). cDNA was synthesized with Superscript IV VILO Mastermix with ezDNase (Thermo Fisher Scientific). qPCR, primer design and validation were conducted as described in Salesse-Smith et al 2024 (61), with the exception that an anneal temperature of 60°C was used for all primer sets. See Table S1 for primer sequences. Primer efficiencies were between 100-105%. Calibrated Normalized Relative Quantities (CNRQ) were calculated using qBase+ software v.3.2 (CellCarta) based on the expression of two reference genes, SbEIF4α and SbActin11.

Total protein was extracted from 1.9 cm^2^ of leaf tissue from the youngest fully expanded leaf of greenhouse-grown sorghum and sugarcane and field-grown sorghum. Protein extraction and western blots were performed as described previously (61). The following primary antibodies were used: anti-HA (AS12 2220, Agrisera) at a 1:5000 dilution, anti-LS (AS03 037, Agrisera) at a 1:10000 dilution, anti-FLAG (F7425-.2MG, Sigma-Aldrich) at a 1:5000 dilution and anti-RAF1 (custom rabbit antibody, Pacific Immunology) at a 1:1000 dilution. For the custom antibody, a 23 amino acid immunograde peptide from the maize RAF1 protein (aa 211-233; KGFPRWRGEEGWEAFSKDSPADC) was synthesized and antisera generated by Pacific Immunology. Relative abundance of LS protein was estimated using Empiria Studio Software version 3.2 (Li-COR, Biosciences).

### Rubisco content, activity and activation measurements

Leaf discs were sampled from the youngest fully expanded leaves of greenhouse-grown sorghum (1.9 cm^2^) and sugarcane (2.5 cm^2^) plants, snap-frozen in liquid nitrogen and stored at −80°C. Samples were taken from the same location gas exchange measurements were made the day after measurements were completed. Soluble protein was extracted in 2 mL of N_2_-sparged extraction buffer: 50 mM EPPS-NaOH pH 8, 1 mM EDTA, 5 mM MgCl_2_, 1% (w/v) PVPP, 5 mM DTT and 1% (v/v) protease inhibitor cocktail (P9599, Sigma-Aldrich). Frozen leaf discs were ground in ice-cold Ten Broeck homogenizers (7 mL, Corning) containing extraction buffer. Total soluble protein was quantified using the Qubit Protein Assay Kit (Q33211, Life Technologies) on the Qubit 3 Fluorometer (Q33216; Life Technologies). Total Rubisco content was measured by ^14^C-CABP binding as previously described (Ruuska et al., 1998; Sharwood et al., 2008). 100 µL of protein lysate was mixed with an equal volume of Rubisco activation buffer (50 mM EPPS-NaOH pH 8, 1 mM EDTA, 20 mM MgCl_2_, 30 mM NaHCO_3_) and let sit at room temperature. After 20 minutes, 60 µM of ^14^C-CABP was added to the mixture and left at room temperature for an additional 30 minutes, and then transferred to ice. 100 µL of the activated Rubisco protein mixture was loaded onto the size-exclusion chromatography columns and fractions containing ^14^C-CABP – Rubisco complexes collected. The amount of ^14^C-CABP was determined by liquid scintillation (Packard Tri-Carb 1900 TR, Canberra Packard Instruments Co). Leaf Rubisco content was calculated as described previously (62).

Rubisco activity was determined at 25°C by incorporation of ^14^CO_2_ into acid-stable products (63). An extended protocol can be found online: dx.doi.org/10.17504/protocols.io.bf8cjrsw. Sorghum (1.7 cm^2^) and sugarcane (2.26 cm^2^) were collected as indicated above. Soluble protein was extracted in 2 mL of N_2_-sparged extraction buffer: 50 mM Bicine-NaOH pH 8.2, 20 mM MgCl_2_, 1 mM EDTA, 2 mM benzamidine, 5 mM ε-aminocaproic acid, 50 mM 2-mercaptoethanol, 10 mM DTT, 1% (v/v) protease inhibitor cocktail, 1 mM phenylmethylsulphonyl fluoride, and 1% (w/v) polyvinylpolypyrrolidone. Supernatant was immediately used to measure initial and total Rubisco activity, with two technical replicates per measurement. Initial activity was measured by adding 25 µl of protein lysate to assay buffer (100 mM Bicine-NaOH pH 8.2, 20 mM MgCl_2_, 10 mM NaH^14^CO_3_ (18.5 kBq μmol^−1^), 2 mM KH_2_PO_4_, and 0.6 mM RuBP). After 30 s reactions were quenched with 100 µl of 10 M formic acid. For total activity measurements, RuBP was added 3 min after the protein lysate to allow for carbamylation of Rubisco and quenched after 30 s. Reactions were dried at 100 °C, then the residue dissolved in 400 µl deionized water and added to scintillation cocktail for acid-stable ^14^C determination by liquid scintillation. Rubisco activation state was calculated as initial/total activity.

### Leaf gas exchange measurements

CO_2_ response (*A*-*C*_i_) and light induction curves were measured on the youngest fully expanded leaves using an infrared gas exchange system LI-6800 fitted with the multiphase flash fluorometer and 6 cm^2^ cuvette (6800-01A, LI-COR Environmental). CO_2_ response curves were measured simultaneously with chlorophyll fluorescence and used to assess *A*, *C*_i_, *g*_sw_, ɸPSII, *V*_c,max_ and *V*_p,max_. *A*-*C*_i_ curves were measured in the field-grown sorghum on July 10^th^, August 4^th^ and August 29^th^. Leaves were acclimated to the following conditions: light intensity of 1800 µmol m^−2^ s^−1^, leaf temperature of 30 °C, CO_2_ reference concentration of 400 µmol mol^−1^ and 65% humidity. Once *A* and *g*_sw_ stabilized, response curves were initiated using the following sequence of reference CO_2_ concentrations: 400, 300, 200, 120, 70, 30, 10, 400, 400, 500, 700, 900, 1200, and 1500 µmol mol^−1^. Measurements were logged 3 to 5 minutes after each new CO_2_ step. *A*-*C*_i_ curves for greenhouse-grown sorghum were measured under the same conditions except for the following changes: a leaf temperature of 28 °C was used and the CO_2_ sequence used concentrations of 600 and 800 µmol mol^−1^ CO_2_ instead of 700 and 900. Sugarcane *A*-*C*_i_ curves used a CO_2_ reference concentration of 450 µmol mol^−1^ and the following sequence of CO_2_: 450, 1500, 1250, 1000, 750, 600, 500, 400, 300, 200, 120, 70, 30, and 10 µmol mol^−1^. *V*_c,max_ and *V*_p,max_ at 25°C were estimated using the *fit_c4_aci* function from the PhotoGEA R package (64), which fits a mechanistic model to the measured CO_2_ response curves (7). In this method, values of *V*_c,max_, *V*_p,max_, and *R*_L_ are varied to find the best fit of the full equation for enzyme-limited assimilation (Equation 4.21 of von Caemmerer 2000) to all points in each curve (29).

Light induction curves were used to assess average *A* in either the first 180 or 300 seconds since the light was turned on and time to 50% *A*_max_, which is determined from the normalized induction curves. Leaves were acclimated to the following conditions: light intensity of 0 µmol m^−2^ s^−1^, leaf temperature of 28°C (greenhouse-grown sorghum) or 30 °C (field-grown sorghum and sugarcane), CO_2_ sample concentration of 400 (sorghum) or 450 µmol mol^−1^ (sugarcane) and 65% humidity. After 30 minutes in the dark, light was increased to 1800 µmol m^−2^ s^−1^ and data points logged every 10 s over the course of 30 min. Average assimilation after 300 seconds of light induction was calculated for sugarcane, while 180 seconds was used for sorghum due to differences in each species overall speed of induction. After gas exchange measurements were completed on the greenhouse-grown sorghum plants, leaf number, tiller number and height were recorded.

### Estimating bundle sheath leakiness using carbon isotope discrimination coupled with leaf gas exchange

A LI-6800 gas exchange system (LI-COR Environmental) with the large light source chamber (6800–03) was coupled to a tunable-diode laser absorption spectroscope (TDLAS model TGA 200A; Campbell Scientific) to measure online carbon isotope discrimination as described previously (24). The TDL was connected to the LI-6800 reference air stream using the port on the back of the sensor head, and connected to the sample air stream using the port on the front of the sensor head. CO_2_-free air (N_2_/O_2_) with 21% O_2_ was created by mixing two gas streams with mass flow controllers (OMEGA Engineering Inc.). This air supplied the gas exchange system, and was used in the calibration to correct for any drift in the TDL during measurements. The TDL was calibrated using a 2-point calibration where a linear fit is applied using the measurements from two calibration tanks with known [^12^CO_2_], [^13^CO_2_] (NOAA Global Monitoring Laboratory). The measurements cycled through four gas streams in the following sequence: calibration points 1500 and 100 ppm CO_2_ (NOAA Global Monitoring Laboratory), LI-COR 6800 reference air stream, leaf chamber air stream, and reference air stream a second time. Each step had a duration of 30 s, except for the leaf chamber air, which had a duration of 600 s for a total cycle time of 720 s. Measurements were averaged over the last 10 s to produce a single data point for the calibration and reference measurements each cycle. For the sample line, measurements were averaged over 10 s intervals, excluding the first 10 s, to have 59 data points each cycle.

The gas exchange system was set to the following conditions: light intensity of 0 µmol m^−2^ s^−1^, leaf temperature of 27°C, flow rate of 300 µmol s^−1^, 21% O_2_, and 800 µmol mol^−1^ CO_2._ Once steady-state respiration was reached, five consecutive measurements were logged to measure dark respiration. Afterwards the system was initiated to auto-log at 10 s intervals over the course of 30 min. Light intensity increased to 1800 µmol m^−2^ s^−1^ upon initiation of the auto program. Sorghum leaves from the field were cut from plants pre-dawn, submerged in water and brought to the lab where they were re-cut underwater and left in the dark until measured. Only steady state measurements of leakiness were calculated (25-30 min after light turned on).

Observed photosynthetic discrimination (Δ^13^C) was derived and bundle sheath leakiness (*ϕ*) was subsequently calculated from gas exchange parameters and Δ^13^C as described previously (65, 66). Data processing and calculations of Δ^13^C and *ϕ* were done using the MATLAB tool developed previously (24), with minor modifications to apply the 2-point calibration discussed above.

### Carbon and nitrogen analysis

Leaf tissue from field grown sorghum was oven dried to a constant weight, ground and total carbon (C), C isotope analysis (δ15C), total nitrogen (N), and N isotope analysis (δ15N) measured using an elemental analyzer (Costech Elemental Combustion System 4010, Costech Analytical Technologies) coupled with a continuous flow isotope ratio mass spectrometer (DeltaV Advantage, Thermo Fisher Scientific). Spinach leaves (National Institute of Standards and technology Standard Reference Material 1570a) and acetanilide (USA Analytical) were used as standards for C and N analysis. Isotope analysis was measured relative to the international standards NBS 19 for C (δ^13^C_V-PDB_ = 1.95*‰)* and atmospheric N_2_ for N (δ^15^N_Air_ = 0‰).

### Statistical analysis

Normality of the data was tested with Shapiro-Wilk’s test, and homoscedasticity with Brown-Forsythe’s test. If criteria for normal distributions and equal variance was met one-way ANOVA followed by Dunnett’s *post hoc* test for transgenic mean comparison against the WT control was performed. Data were considered significant at *P* < 0.05, and at *P* < 0.1 when specified. If criteria for normality was violated Wilcoxon’s non-parametric test was applied. Analysis of field growth traits (Fig. 4, Table S2) was performed using a randomized block design with 6 blocks. Statistical tests used are indicated in each figure or table legend. Jmp pro version 17.0.0 software was used for all statistical analyses.

## Acknowledgements

We thank Montgomery Flack for maintenance of the greenhouse plants. We thank David Drag, Ben Thompson, and Andy Wszalek for help with the maintenance of the sorghum field experiment and field harvest. We thank Edward Lochocki for assistance during field work and Samantha Stutz for maintenance of the TDL. We also thank Mike Masters and Chuck Hyde for performing the C and N analysis and Ben Haas for initial help with plasmid assembly. We are grateful to Dr. Berkley Walker for the kind gift of RuBP. This work was funded by the DOE Center for Advanced Bioenergy and Bioproducts Innovation (U.S. Department of Energy, Office of Science, Biological and Environmental Research Program under Award Number DE-SC0018420). Any opinions, findings, and conclusions or recommendations expressed in this publication are those of the author(s) and do not necessarily reflect the views of the U.S. Department of Energy.

## Author Contributions

CES and SPL conceived of and designed the experiments. TEC coordinated sorghum transformation, MG assembled the vector and ZG generated and confirmed the transgenic sorghum lines. WW performed sorghum copy number analysis. FA coordinated sugarcane transformation, BK and TN, generated and confirmed the transgenic sugarcane lines. NA and CES assembled the recombinant DNA vector for sugarcane, and performed experimental measurements. CES analyzed the data. CES and SPL wrote the manuscript with contributions and review from all authors.

## Supporting Information

Fig. S1: Screening T1 transgenic sorghum lines.

Fig. S2: Screening T0 transgenic sugarcane lines.

Fig. S3: Gene expression in greenhouse and field grown sorghum.

Fig. S4: Normalized induction curves from greenhouse and field grown sorghum.

Fig. S5: Plant growth traits in greenhouse grown sorghum plants.

Fig. S6: Sorghum field experimental design and weather conditions.

Fig. S7: Steady-state BS leakiness of field grown sorghum.

Fig. S8: Normalized induction curves from greenhouse-grown sugarcane.

Fig. S9: Plant growth traits in greenhouse grown sugarcane plants.

Fig. S10: Average yield of sorghum and sugarcane over the last 20 to 30 years.

Fig. S11: Light intensity directly outside the greenhouse.

Table S1: List of primers used in this manuscript.

Table S2: Summary of harvest measurements and yield characterization of field-grown sorghum.

Table S3: Summary of leaf mass per area (LMA), chlorophyll content (SPAD value), leaf carbon and nitrogen content and total soluble protein (TSP) of sorghum and sugarcane plants.

