## Supplemental Figures 1-11, Tables 1-3 for "Adapting C_4_ photosynthesis to atmospheric change and increasing productivity by elevating Rubisco content in Sorghum and Sugarcane"

[1]Carl R. Woese Institute for Genomic Biology, University of Illinois at Urbana-Champaign, Urbana, IL, USA; [2]Departments of Plant Biology and of Crop Sciences, University of Illinois at Urbana-Champaign, Urbana, IL, USA; [3]DOE Center for Advanced Bioenergy and Bioproducts Innovation, Urbana-Champaign, IL, USA; [4]Agronomy Department, Plant Molecular and Cellular Biology Program, Genetics Institute, University of Florida, IFAS, Gainesville, Florida, USA; [5]DOE Center for Advanced Bioenergy and Bioproducts Innovation, Gainesville, FL, USA; [6]Department of Agronomy and Horticulture, University of Nebraska-Lincoln, Lincoln, NE, USA; [7]DOE Center for Advanced Bioenergy and Bioproducts Innovation, Lincoln, NE, USA

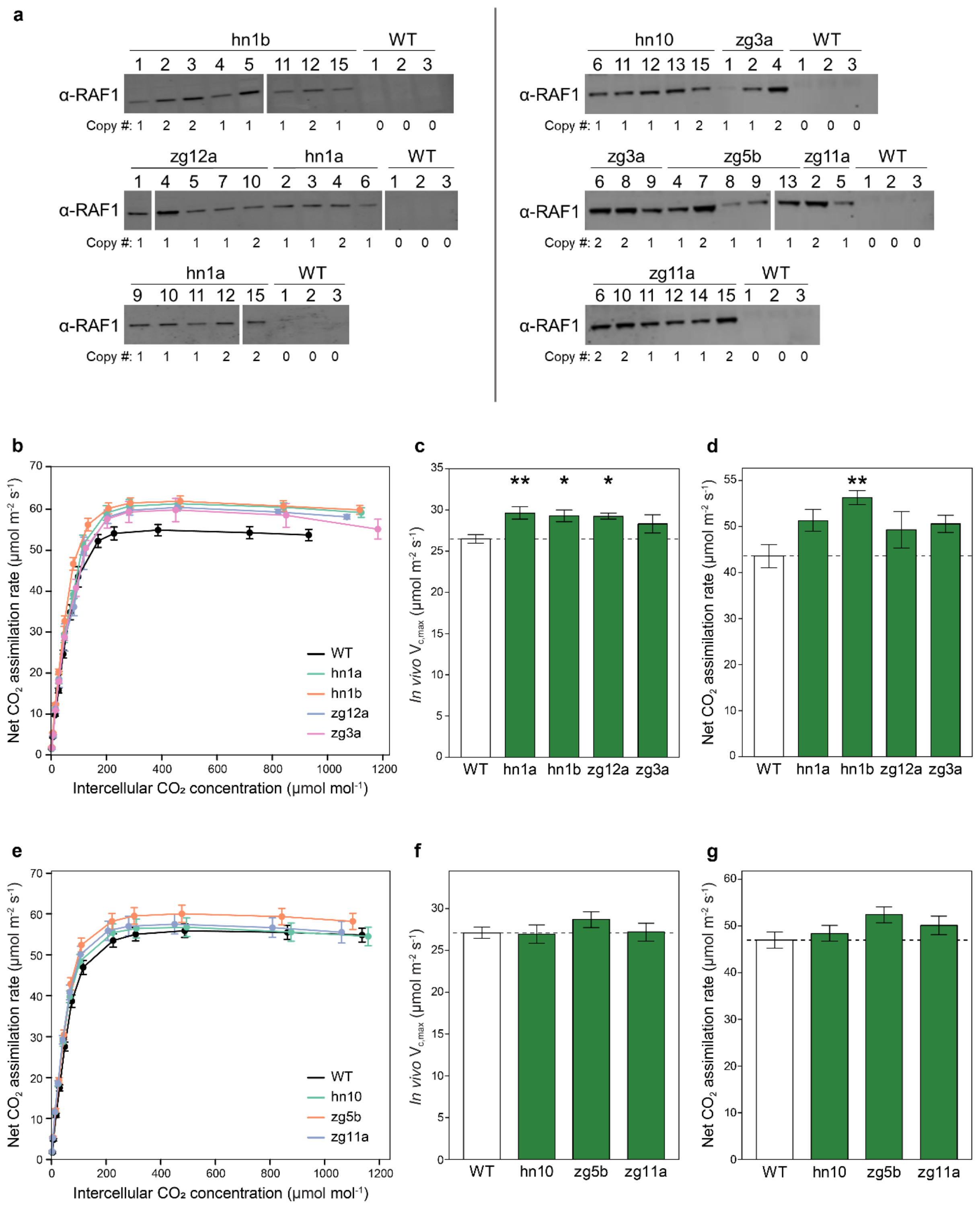


**Fig. S1: Screening T1 transgenic sorghum lines.**

**(a)** Total soluble protein isolated on a leaf area basis from sorghum, analyzed by immunoblot and probed with anti-RAF1 antibody. Copy number of NPTII obtained by digital droplet PCR is indicated below each sample. **(b and e)** Response of net $CO_2$ assimilation ($A_{sat}$) to intercellular [CO2]. **(c and f)** Maximum *in vivo* Rubisco carboxylation rate ($V_{c,max}$) at 25 °C estimated from response curves. **(d and g)** $A_{sat}$ measured at 400 μmol mol$^{-1}$ $CO_2$. Values are shown as the mean ± SEM. Asterisks indicate significant differences between WT and the transgenic line (**P < 0.05, *P<0.1); one-way ANOVA, Dunnett's post hoc test. For gas exchange screening the seven transgenic events were split into two sets. Set 1 (b-d) n=4. Set 2 (e-f) n=7-8.

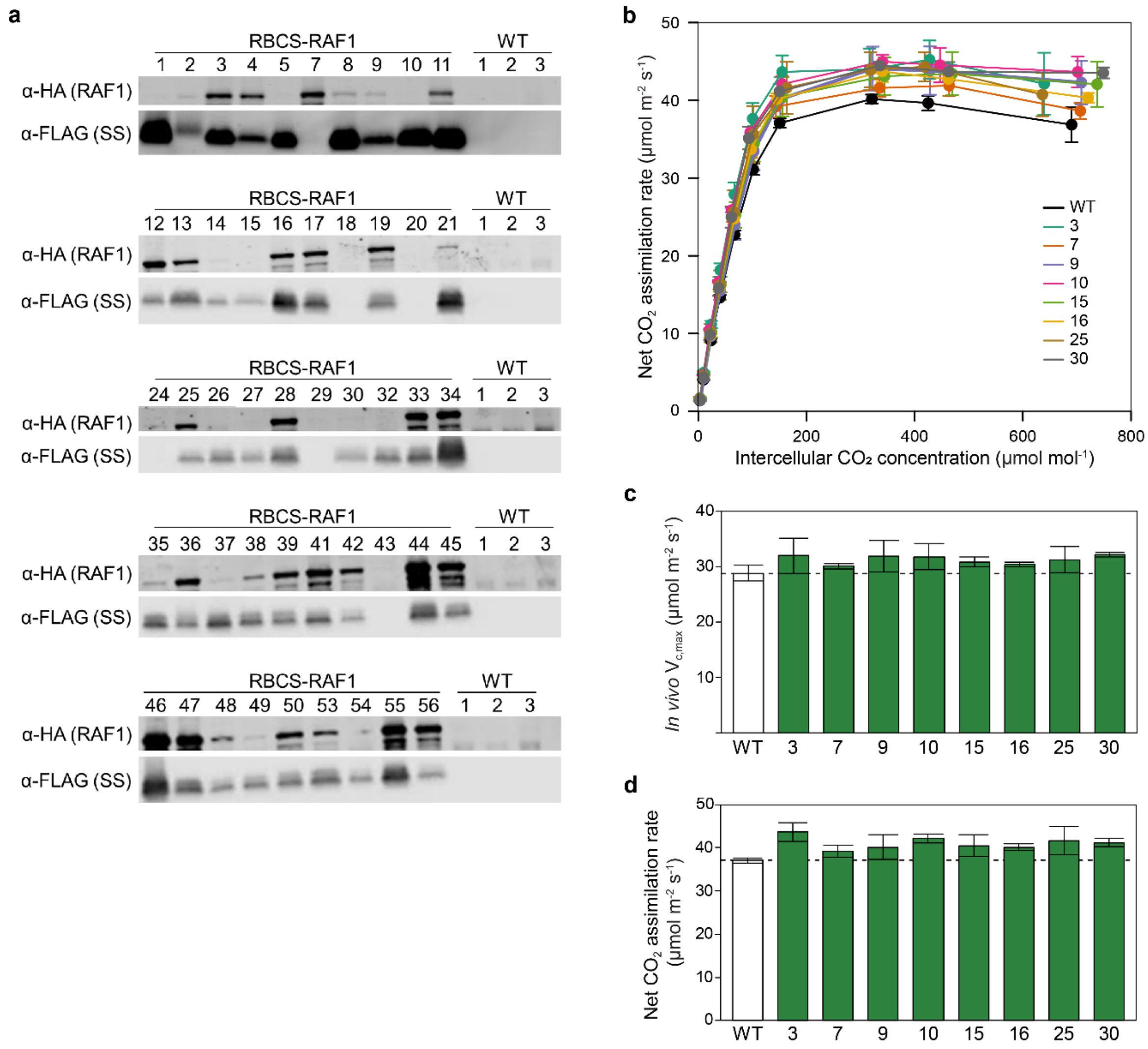


**Fig. S2: Screening T0 transgenic sugarcane lines.**

**(a)** Total soluble protein isolated on a leaf area basis from sugarcane, analyzed by immunoblot and probed with antibodies indicated to the left. **(b)** Response of net $CO_2$ assimilation ($A_{sat}$) to intercellular [CO2]. **(c)** Maximum *in vivo* Rubisco carboxylation rate ($V_{c,max}$) at 25 °C estimated from response curves (d) $A_{sat}$ measured at 400 µmol mol$^{-1}$ $CO_2$. Values are shown as the mean ± SEM (n=3 technical replicates from different leaves).

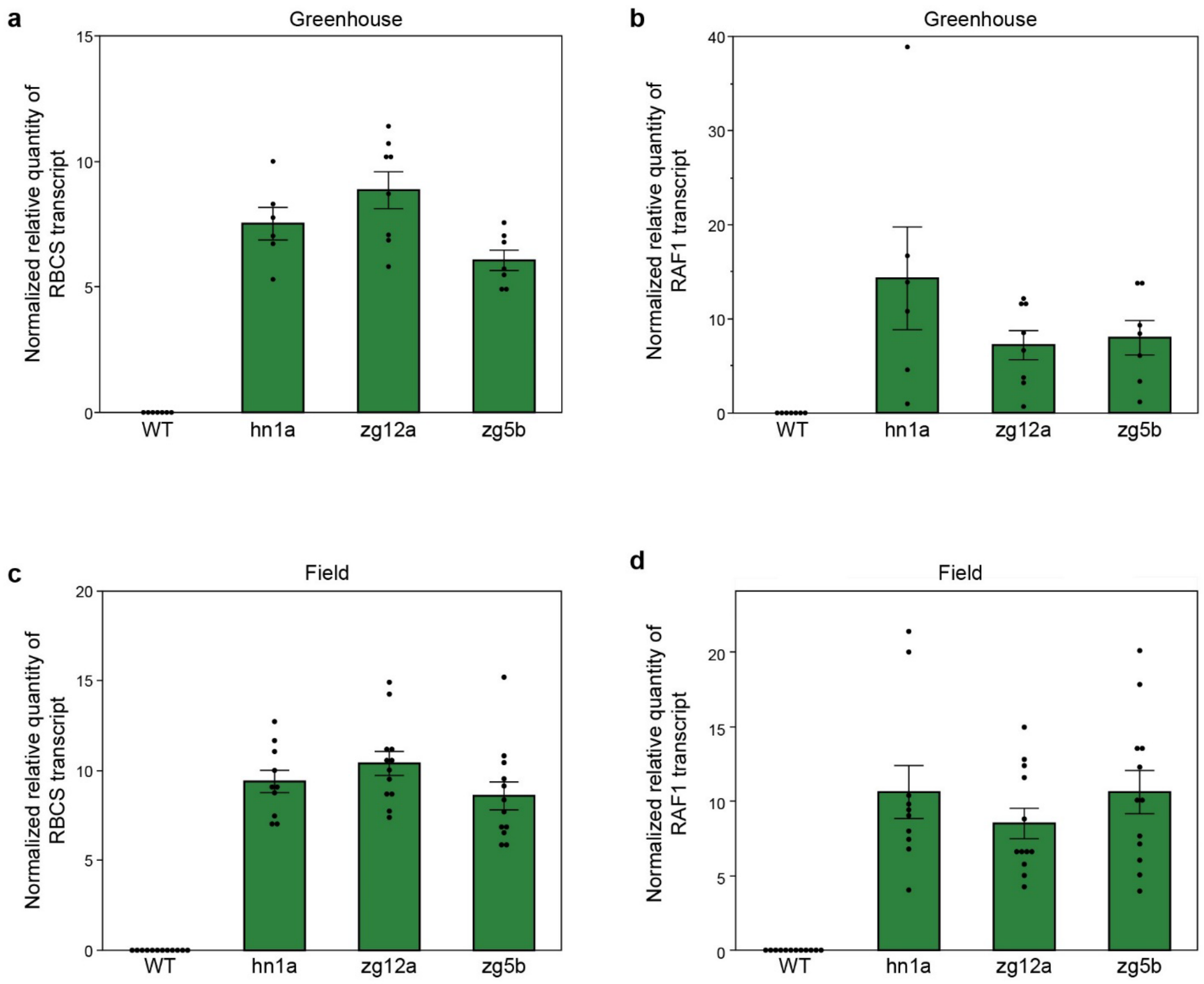


**Fig. S3: Gene expression in greenhouse and field grown sorghum.**

**(a)** qPCR analysis of *ZmRbcS* and **(b)** *ZmRaf1* gene expression in three independent transgenic sorghum events grown in the greenhouse (n = 6-8). **(c)** *ZmRbcS* and **(d)** *ZmRaf1* gene expression in field grown sorghum (n = 10-12). Values are shown as the mean ± SEM.

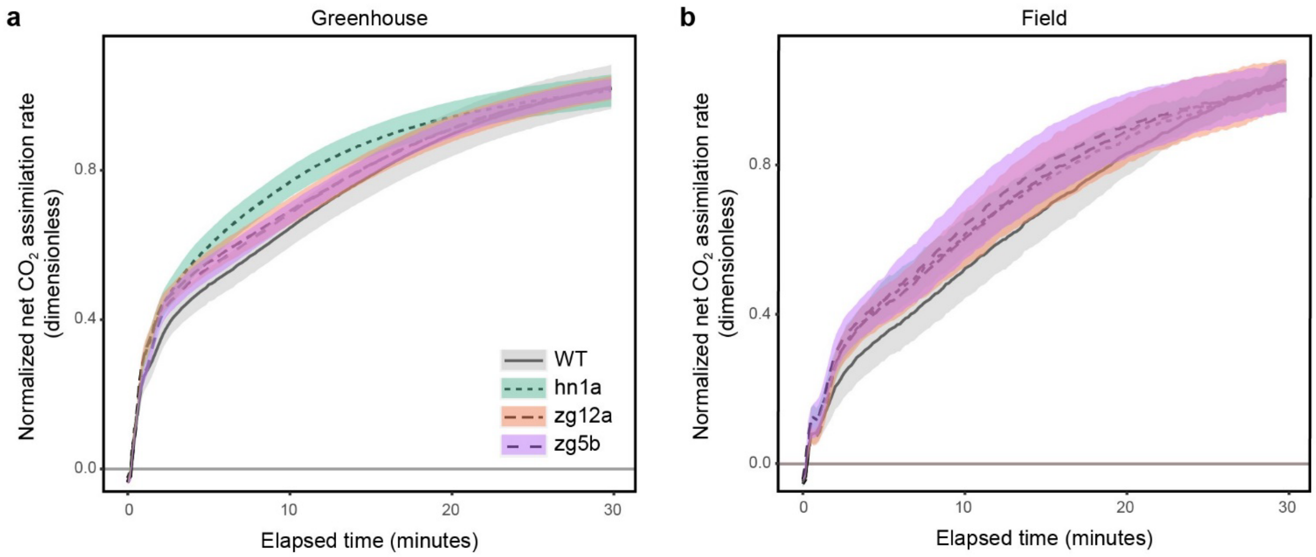


**Fig. S4: Normalized induction curves from greenhouse and field grown sorghum.**

**(a)** Normalized induction of net $CO_2$ assimilation during the first 30 min of illumination at PPFD 1800 µmol $m^{-2}$ $s^{-1}$ in the greenhouse (n = 8-9) and **(b)** in field grown sorghum (n = 4-5). Values are shown as the mean ± SEM and were normalized by the average value over the last 5 minutes of illumination from Fig. 2e and 3b respectively.

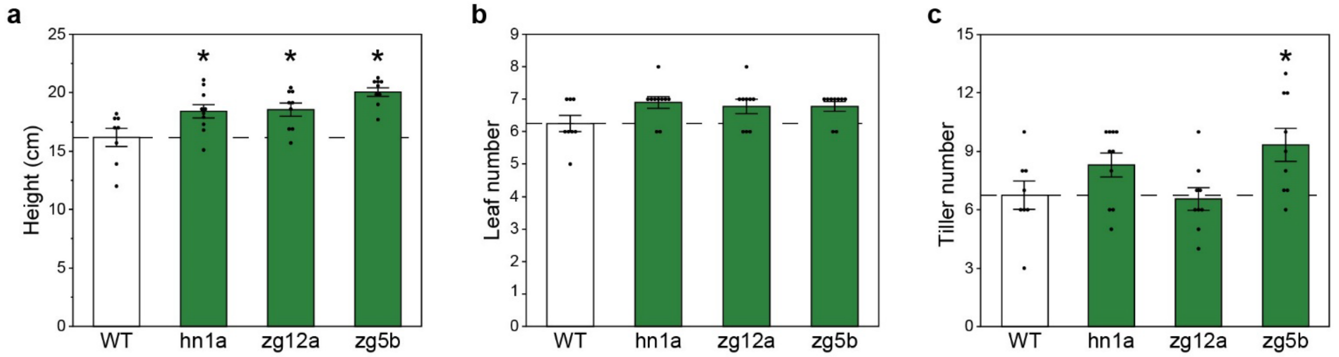


**Fig. S5: Plant growth traits in greenhouse grown sorghum plants.**

**(a)** Plant height, **(b)** leaf number, **(c)** tiller number. Values are shown as the mean ± SEM (n = 8-10). Asterisks indicate significant differences between WT and the transgenic line (*P < 0.05); one-way ANOVA, Dunnett's post hoc test.

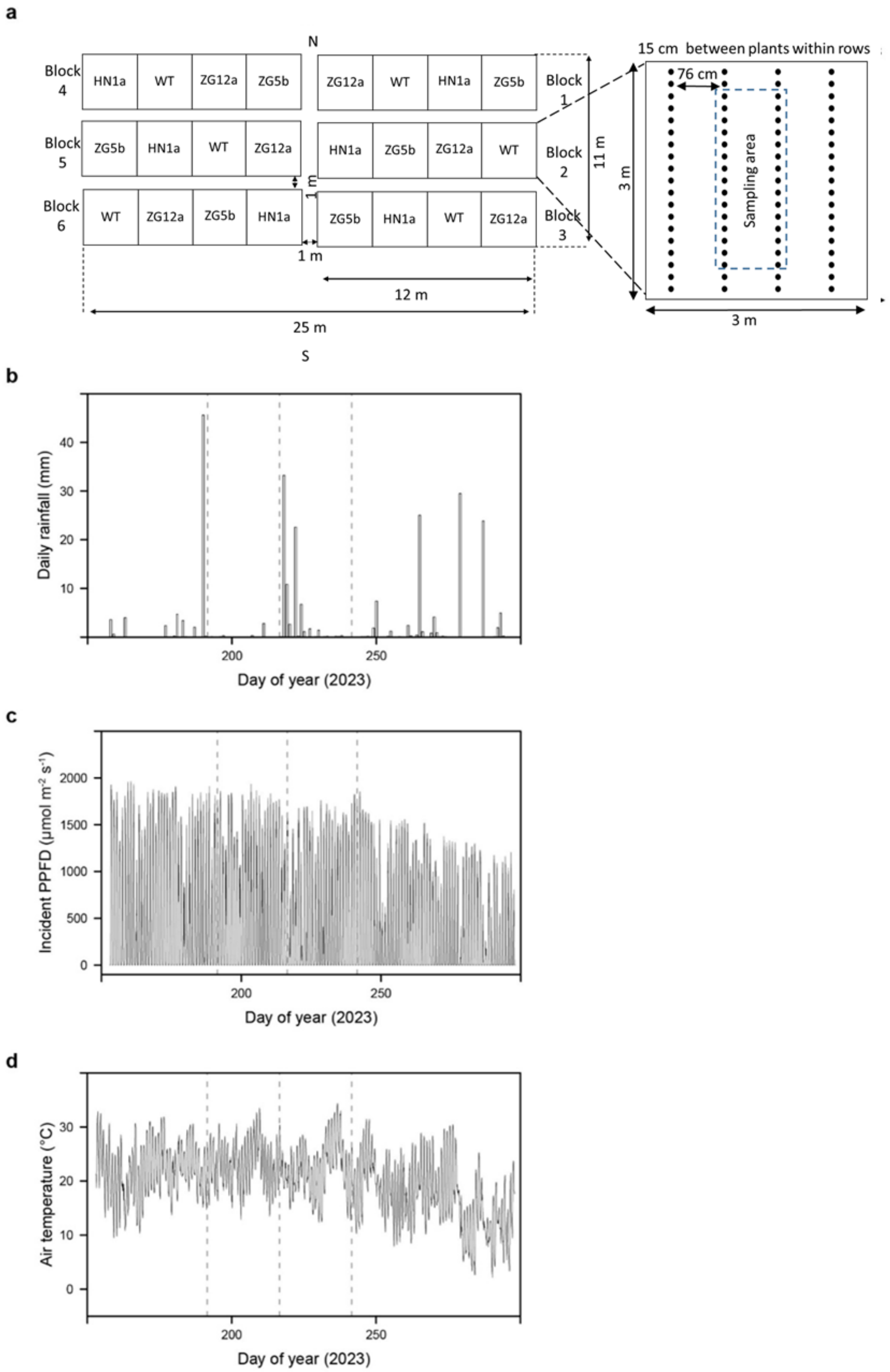


**Fig. S6: Sorghum field experimental design and weather conditions.**

**(a)** Schematic of field experimental set up. A randomized block design was used. Genotypes were randomly assigned to one of four positions in each of the six blocks; ● on the inset shows the location of individual plants. **(b)** Total daily rainfall, **(c)** light intensity and **(d)** air temperature from June 2$^{nd}$ (DOY 153 - date sorghum sowed) until October 4$^{th}$, 2023 (DOY 297 - date sorghum harvested). (c-d) Data are shown as averages over the preceding hour. Dashed vertical lines show dates of gas exchange measurements; July 10$^{th}$ (DOY 191), August 4$^{th}$ (DOY 216), and August 29$^{th}$ (DOY 241).

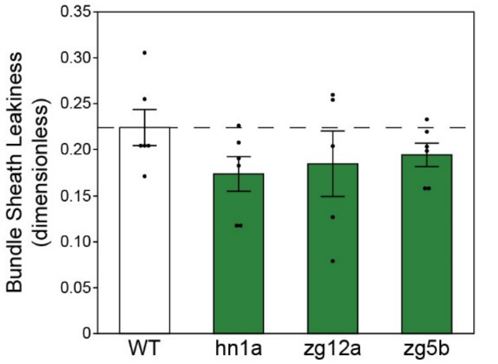


**Fig. S7: Steady-state BS leakiness of field grown sorghum.**

Bundle-sheath leakiness measurements. Values are shown as the mean ± SEM (n = 5-6). No significant differences, one-way ANOVA, Dunnett's post hoc test.

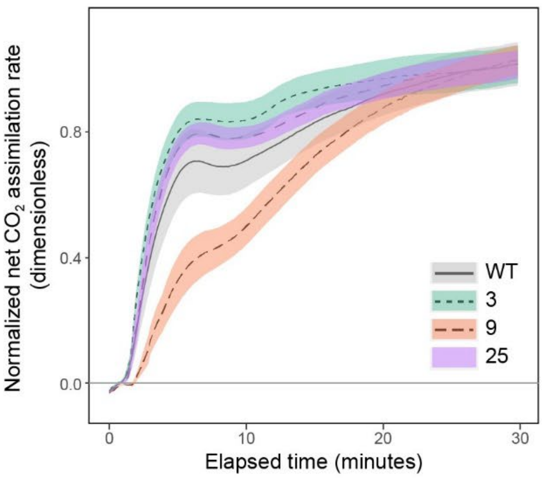


**Fig. S8: Normalized induction curves from greenhouse-grown sugarcane.**

**(a)** Normalized induction of net $CO_2$ assimilation during the first 30 min of illumination at PPFD 1800 µmol $m^{-2}$ $s^{-1}$ (n = 7-8). Values are shown as the mean ± SEM and were normalized by the average value over the last 5 minutes of illumination from Fig. 5b.

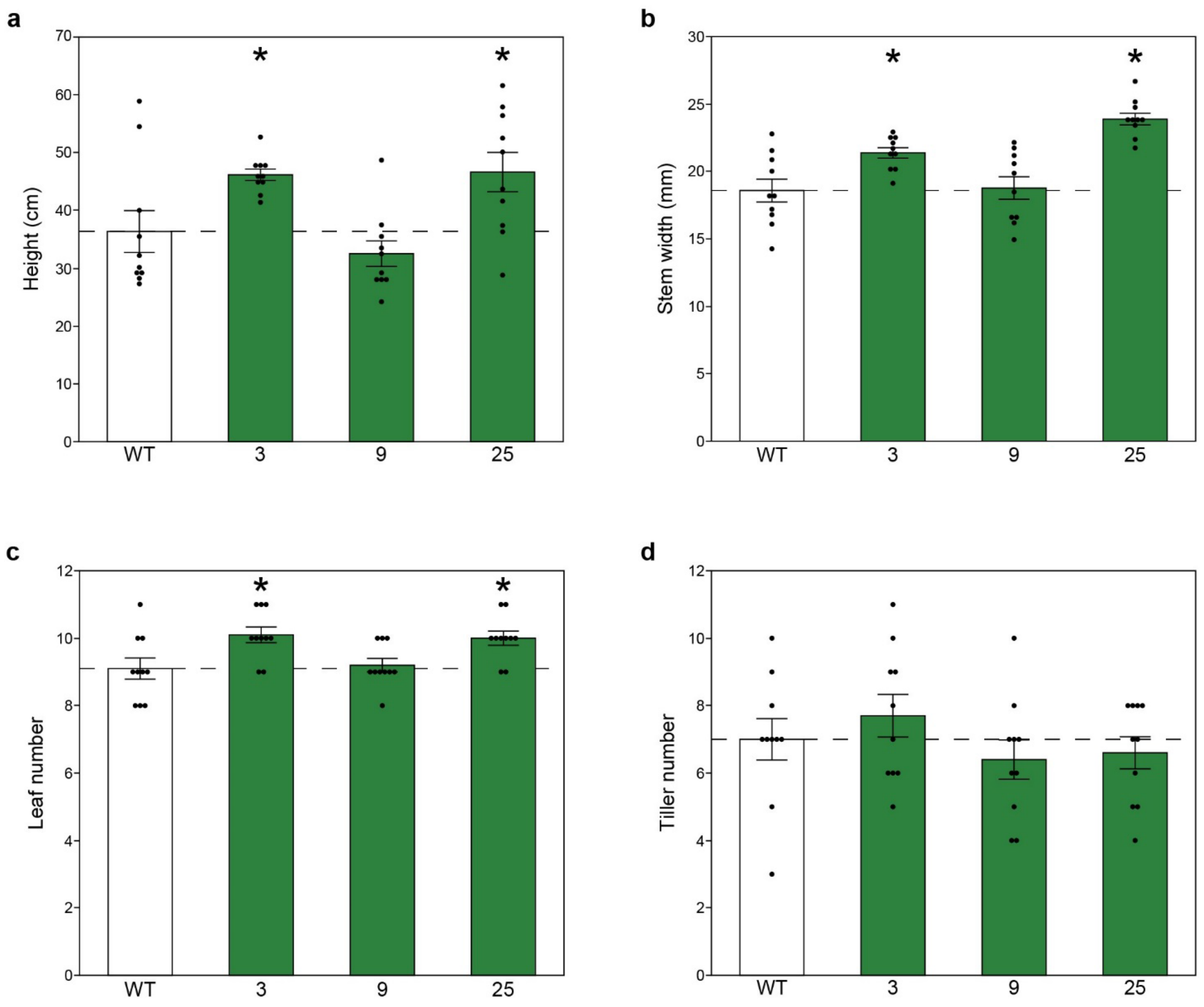


**Fig. S9: Plant growth traits in greenhouse grown sugarcane plants.**

**(a)** Plant height, **(b)** stem width, **(c)** leaf number and **(d)** tiller number. Values are shown as the mean $\pm$ SEM (n = 10). Asterisks indicate significant differences between WT and the transgenic line (*P < 0.05); one-way ANOVA, Dunnett's post hoc test.

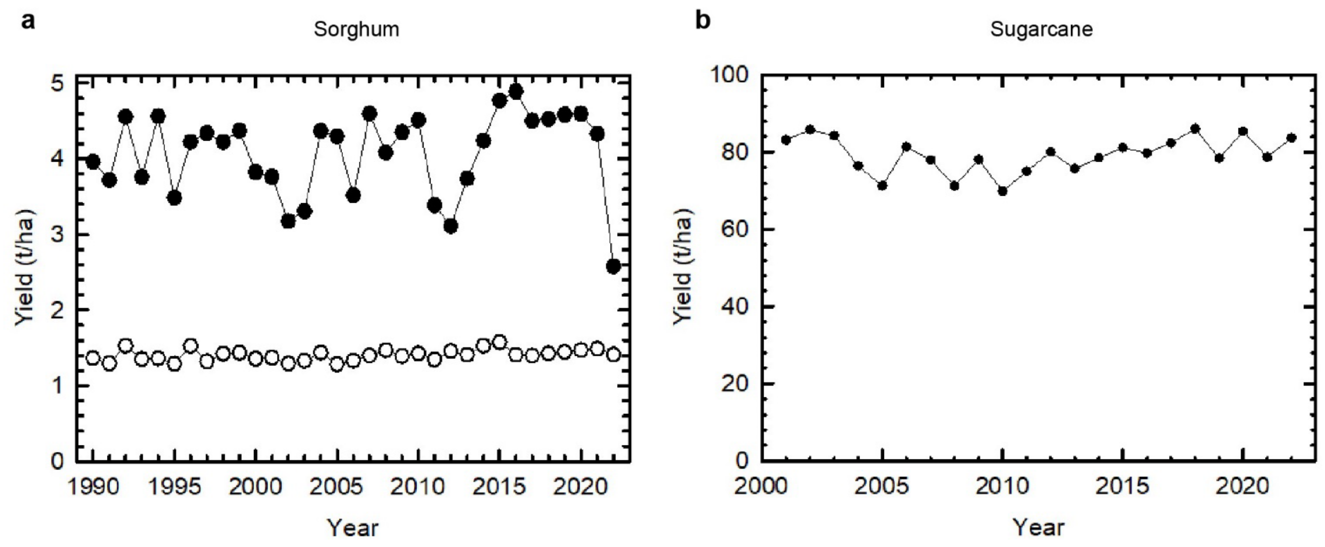


**Fig. S10: Average yield of sorghum and sugarcane over the last 20 to 30 years.**

**(a)** Official figures from FAOstat (2024) for average World (open symbols) and USA (closed symbols) sorghum grain yields for 1990-2022, and **(b)** yield of harvested sugarcane stalks in the USA from 2001-2022.

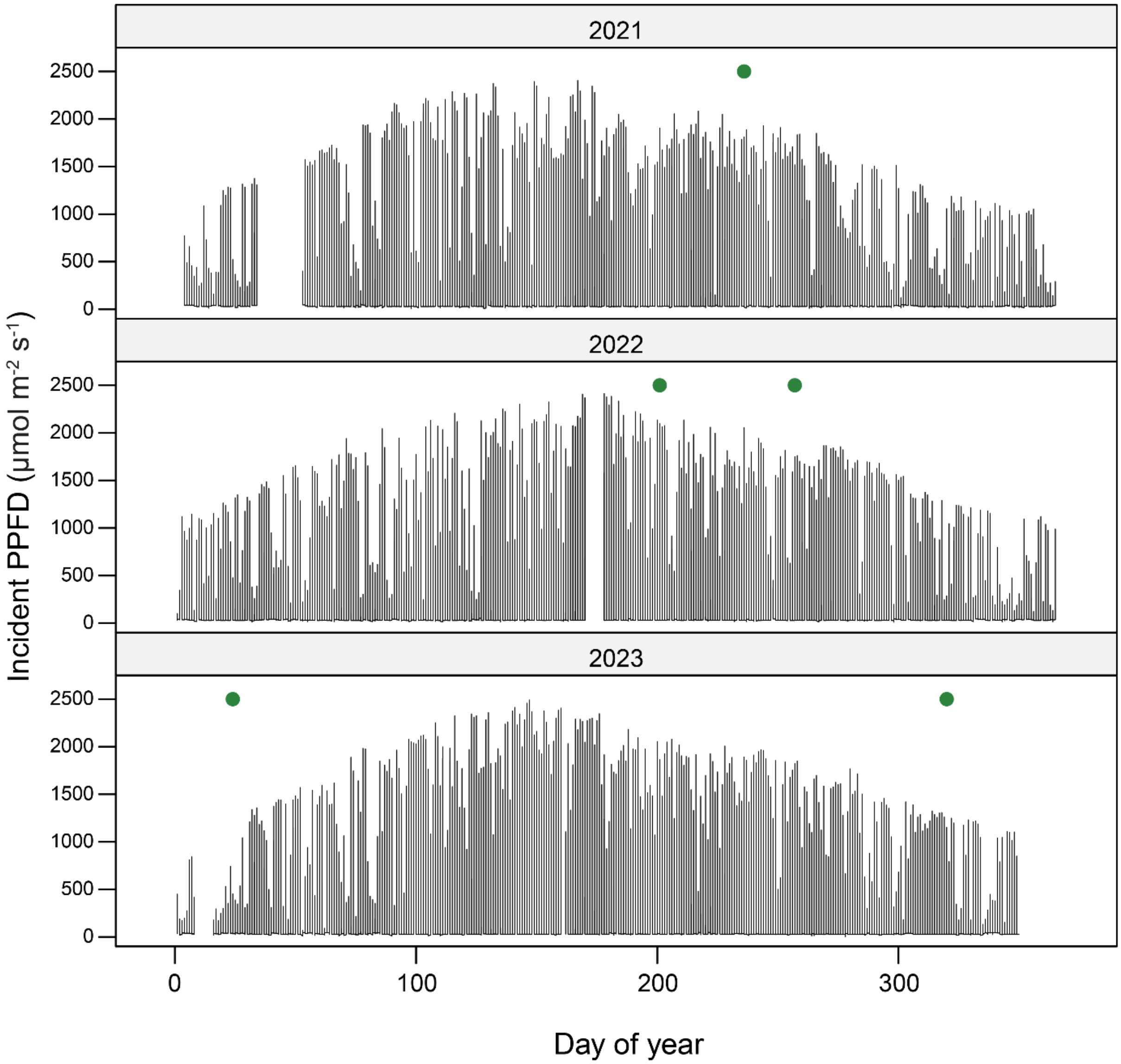


**Fig. S11: Light intensity directly outside the greenhouse.**

Outside irradiance data for greenhouse experiments presented in this manuscript. Filled green circles show dates for gas exchange measurements: sugarcane screening on August 24-27, 2021 (DOY 236-239); sorghum screening (set I) on July 20, 2022 (DOY 201); sorghum screening (set II) on September 14-15, 2022 (DOY 257-258); main sorghum experiment on January 24-26, 2023 (DOY 24-26); main sugarcane experiment on November 16-17, 2023 (DOY 320-321).

**Table S1. List of primers used in this manuscript. * designates reference genes.**

| Primer name | Sequence | Type | Amplicon length (bp) |
|---|---|---|---|
| ddNPTII_F | TACGCTTGATCCGGCTAC | Sb ddPCR | 77 |
| ddNPTII_R | CTTCCATCCGAGTACGTG | Sb ddPCR | |
| *Sb ENOL-2_F | TGAGGACCCTTTTGATCAGG | Sb ddPCR | 135 |
| *Sb ENOL-2_R | CAAGCCTTCTTGCCAATAGC | Sb ddPCR | |
| So Construct_F | TAATGGTGGTGTAGCTCGACG | So PCR | 209 |
| So Construct_R | TGGCTTGGTTAGGTTTGGCT | So PCR | |
| *NPTII_F | TACCTGCCCATTCGACCACC | So PCR | 345 |
| *NPTII_R | TAAAGCACGAGGAAGCGGTC | So PCR | |
| ZmRBCS_F | GCGTCTCGAACTTCTTGTTGC | Sb qPCR | 111 |
| ZmRBCS_R | TGGACTGATGTGTGTTGCCC | Sb qPCR | |
| ZmRAF1_F | CCAAGGAGTCGAAGTAGGGAA | Sb qPCR | 149 |
| ZmRAF1_R | CTTCAACCCGTCTTCCATCCA | Sb qPCR | |
| *SbEIF4α_F | CAACTTTGTCACCCGCGATGA | Sb qPCR | 144 |
| *SbEIF4α_R | TCCAGAAACCTTAGCAGCCCA | Sb qPCR | |
| *SbActin11_F | AACTGCAGATGTGGATTGCCAAGG | Sb qPCR | 115 |
| *SbActin11_R | ATAATGGCTCCTCTCGGCTTGCAT | Sb qPCR | |

**Table S2. Summary of harvest measurements and yield characterization of field-grown sorghum.** Values are shown as the mean ± SEM. Values in bold are significantly different from WT at $P < 0.05$.

| Parameter | WT | RBCS-RAF1 Hn1a | RBCS-RAF1 Zg12a | RBCS-RAF1 Zg5b | Sample number (*N*) |
|---|---|---|---|---|---|
| **Sorghum - Field** | | | | | |
| Panicle dry weight, with seed (kg) | 3.53 ± 0.15 | 2.99 ± 0.23 | 3.24 ± 0.12 | 3.24 ± 0.25 | 6 plots |
| Panicle dry weight, without seed (kg) | 0.654 ± 0.030 | 0.657 ± 0.046 | 0.631 ± 0.023 | 0.648 ± 0.050 | 6 plots |
| Above ground biomass (kg) | 8.10 ± 0.32 | 8.24 ± 0.53 | 8.34 ± 0.30 | 8.75 ± 0.66 | 6 plots |
| Number of panicles per plot | 69.5 ± 2.36 | 70.7 ± 4.14 | 56.7 ± 3.28 | 60.8 ± 4.21 | 6 plots |
| 100 seed weight (g) | 4.02 ± 0.050 | **3.78 ± 0.029** | **3.83 ± 0.033** | 4.06 ± 0.018 | 6 plots |
| Average seed weight per panicle (g) | 41.3 ± 1.03 | **32.8 ± 1.19** | 46.6 ± 2.81 | 42.5 ± 1.09 | 6 plots |
| Panicle emergence (DOY) | 227 ± 0.48 | **233 ± 0.34** | **230 ± 0.40** | **232 ± 0.54** | 6 plots |

**Table S3. Summary of leaf mass per area (LMA), chlorophyll content (SPAD value), leaf carbon and nitrogen content and total soluble protein (TSP) of sorghum and sugarcane plants.** (a) Relative abundance of LS protein estimated from Fig. 1b (sorghum) and Fig. 1c (sugarcane). Values are shown as the mean ± SEM. Values in bold are significantly different from WT at $P < 0.05$.

| Parameter | WT | RBCS-RAF1 Hn1a | RBCS-RAF1 Zg12a | RBCS-RAF1 Zg5b | Sample number (*N*) |
|---|---|---|---|---|---|
| **Sorghum - Greenhouse** | | | | | |
| TSP (µg/ml) | 289.02 ± 27.96 | 280.22 ± 25.68 | 300.22 ± 30.36 | 286.58 ± 25.09 | 8 |
| Relative abundance[a] of LS protein | 1 ± 0.35 | 1.49 ± 0.17 | 1.35 ± 0.30 | 1.30 ± 0.23 | 3 |
| **Sorghum - Field** | | | | | |
| LMA (g/m$^2$) | 55.18 ± 1.18 | 56.21 ± 0.84 | 54.80 ± 0.94 | 55.33 ± 0.80 | 12 |
| SPAD | 55.30 ± 2.47 | 51.52 ± 2.28 | 56.57 ± 2.35 | 52.28 ± 0.756 | 15 |
| Leaf Carbon (%) | 43.17 ± 0.32 | 42.91 ± 0.51 | 42.16 ± 0.61 | 42.67 ± 0.45 | 11-12 |
| $\delta^{13}$C (‰) | -12.17 ± 0.018 | -12.23 ± 0.043 | -12.18 ± 0.030 | **-12.26 ± 0.021** | 10-12 |
| Leaf Nitrogen (%) | 3.14 ± 0.081 | 3.21 ± 0.085 | 3.21 ± 0.060 | 3.18 ± 0.058 | 12 |
| $\delta^{15}$N (‰) | 4.76 ± 0.21 | 4.68 ± 0.094 | 4.24 ± 0.14 | 4.92 ± 0.18 | 11-12 |
| Parameter | WT | RBCS-RAF1 3 | RBCS-RAF1 9 | RBCS-RAF1 25 | Sample number (*N*) |
| **Sugarcane - Greenhouse** | | | | | |
| LMA (g/m$^2$) | 47.53 ± 2.99 | 42.19 ± 0.85 | 45.31 ± 1.32 | 41.50 ± 1.11 | 4-5 |
| SPAD | 55.06 ± 1.22 | 55.77 ± 0.83 | 58.00 ± 1.11 | 54.49 ± 1.10 | 10 |
| TSP (µg/ml) | 244.67 ± 9.28 | 245.67 ± 4.74 | 244.67 ± 8.65 | 246.67 ± 5.88 | 6 |
| Relative abundance[a] of LS protein | 1 ± 0.079 | **1.20 ± 0.055** | 0.91 ± 0.079 | 1.10 ± 0.038 | 3 |
